# Learning attentional templates for value-based decision-making

**DOI:** 10.1101/2023.07.25.550426

**Authors:** Caroline I. Jahn, Nikola T. Markov, Britney Morea, R. Becket Ebitz, Timothy J. Buschman

**Author notes:** Corresponding authors: Timothy J. Buschman, and Caroline I. Jahn,. Lead contact: Timothy J. Buschman,.

## Abstract

Attention filters sensory inputs to enhance task-relevant information. It is guided by an ‘attentional template’ that represents the stimulus features that are relevant for the current task. To understand how the brain learns and uses new templates, we trained monkeys to perform a visual search task that required them to repeatedly learn new attentional templates. Neural recordings found templates were represented across prefrontal and parietal cortex in a structured manner, such that perceptually neighboring templates had similar neural representations. When the task changed, a new attentional template was learned by incrementally shifting the template towards rewarded features. Finally, we found attentional templates transformed stimulus features into a common value representation that allowed the same decision-making mechanisms to deploy attention, regardless of the identity of the template. Altogether, our results provide new insight into the neural mechanisms by which the brain learns to control attention and how attention can be flexibly deployed across tasks.

## Introduction

Cognitive control focuses on the stimuli, thoughts, and actions that are relevant to the current task. In particular, feature-based visual attention selects those stimuli that have task-relevant features. This is an everyday experience: when hailing a taxicab in New York City, we might attend to yellow stimuli. Feature-based attention is guided by an ‘attentional template’ that contains the set of stimulus features that are relevant for the current task^1–5^. Previous work has shown attentional templates are represented in prefrontal and parietal cortex^6–8^. Both regions are active when attention shifts toward specific features^9–11^ and lesioning (or inactivating) these regions impairs a subject’s ability to attend to features^9,12–16^.

As the task changes, so does the information that is relevant. So, the brain must flexibly adapt, continuously learning and re-learning the attentional template that is optimal for each task. For example, while yellow may be a good attentional template for finding taxis in New York City, the template must be updated to beige in Berlin and red in Costa Rica. The neural mechanisms for learning attentional templates are largely unknown. One hypothesis is that templates are learned through reinforcement-learning. Consistent with this, psychophysical studies have shown that reward can be used to learn new attentional templates^17–23^. Furthermore, rewarded stimuli attract attention^24,25^ and many value-learning studies are thought to also measure changes in attention toward rewarded stimuli^26^. This makes sense; when learning a new task, rewarded stimuli or features are likely to be task-relevant and should be attended. However, most neurophysiological studies of attention provide the subjects with the template and so we do not yet know how rewards shape attentional templates during learning.

Once learned, the attentional template influences sensory processing in order to select task-relevant stimuli^8^. This is thought to occur in a top-down manner, with templates in prefrontal and parietal cortex biasing sensory representations across the brain^27–34^. By strengthening the neural representation of task-relevant features, attentional templates can improve discrimination and influence decisions^35,36^. While these results have provided foundational insight into attention, it remains unclear how attention rapidly reconfigures sensory processing and decision-making as the attentional template flexibly changes between tasks.

To better understand how attentional templates are learned, and how these templates flexibly bias decision-making, we trained monkeys to perform a visual search task that required repeatedly learned new attentional templates in a continuous stimulus space (color). Neural recordings in prefrontal and parietal cortex found attentional templates were represented in both regions. Template representations were structured, such that perceptually similar templates had similar neural representations. Furthermore, we found that when the behavioral task changed, the attentional template was learned incrementally: on every trial, the neural representation of the template shifted towards features that were rewarded. Finally, we found the attentional template transformed stimulus representations into a generalized value representation, allowing the animal to attend to the target stimulus, regardless of the stimulus’ color or the current attentional template. Altogether, our results provide new insight into the neural mechanisms by which the control of attention can adaptively be learned and flexibly deployed across tasks.

## Results

We trained two monkeys to perform a visual search task where they repeatedly learned new attentional templates. On each trial, the monkeys used featural attention to search a visual array of three randomly colored stimuli (Fig. 1A-B; see methods for details). The monkey selected one stimulus from the array by making an eye movement to it, and then received the associated reward. The amount of reward associated with each stimulus was inversely proportional to the angular distance between the color of the stimulus and the template color (Fig. 1A, right; smaller distance along the color wheel equaled greater reward). Therefore, on each trial, the animals used their attentional template to search the array of stimuli in order to identify, and select, the stimulus with the ‘best’ color while ignoring the two distracting stimuli. Overall, the monkeys performed the task well: monkey B (and S) selected the best available stimulus on 65% (67%) of trials.

**Fig. 1.**
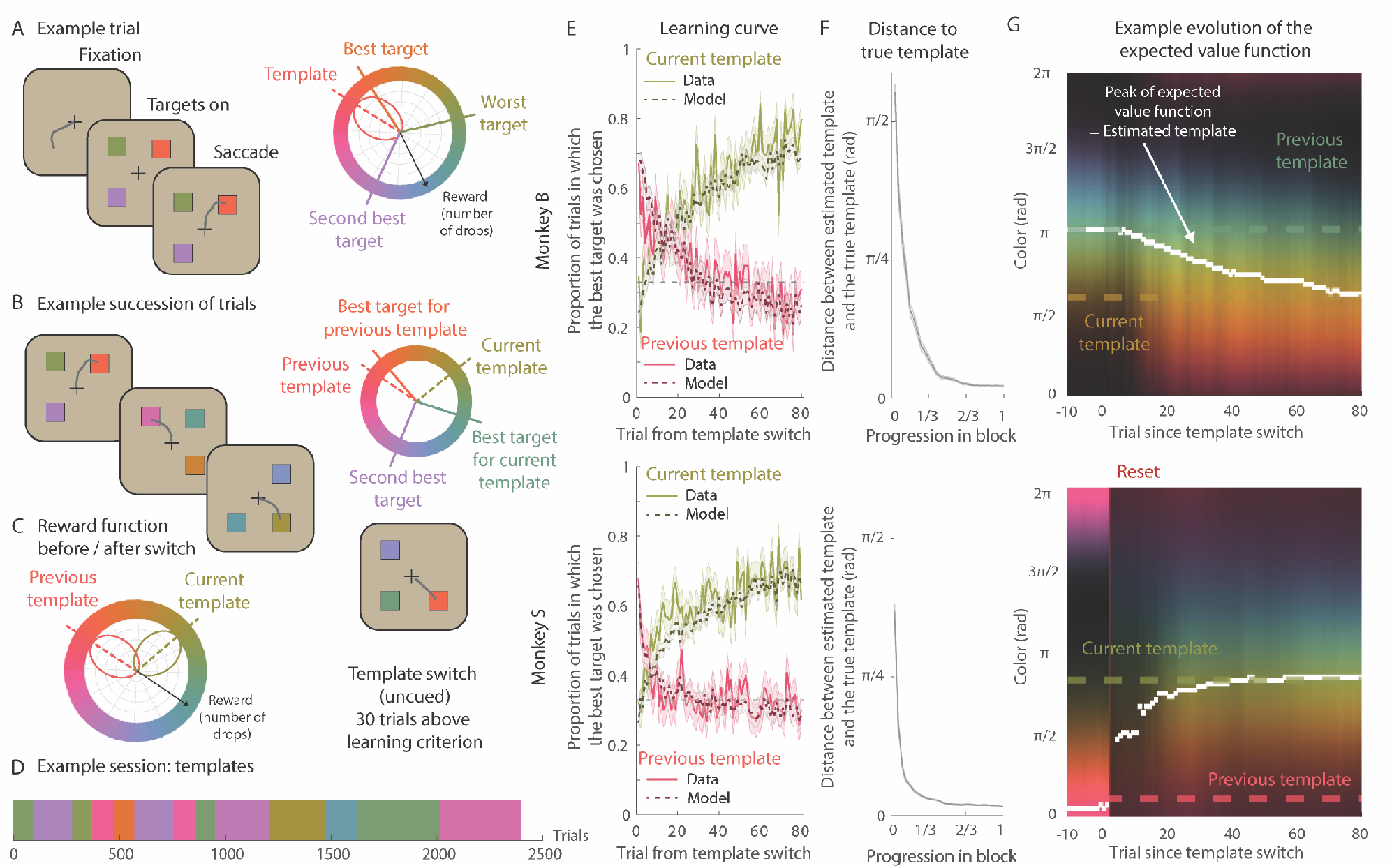
Attentional template learning task and behavior. (**A**) Example trial. A trial started with monkeys fixating a central dot. Three targets of randomly assigned colors appeared. A target was chosen by 50ms fixation. On the color wheel, the reward function (in pink, centered on the (true) template color, the color with the highest reward) defines the amount of reward each target color yields (number of liquid drops) across the color wheel. The target closest to the template color in the color wheel yields the most reward (here, the template color is pink, so the best target is orange). (**B**) Example succession of trials. The targets’ colors and locations (4 possible locations) were random and changed on each trial such that monkeys could not predict either. After monkeys reached a learning criterion (selecting the best target in 80 or 85% of trials in 30 consecutive trials), the template color changed. This switch was uncued and so monkeys would initially choose the best target according to the previous template (here orange) before relearning. (**C**) Reward function before and after the template switch. When the template color changes, by definition, the most rewarded color changes and so does the reward function over color wheel. Before the template switch, the reward function (in pink) was centered on the previous template, such that colors closest to it yielded the most reward. After the template switch, the reward function (in green) was centered on the current template, changing the amount of reward associated with each color. (**D**) Sequence of templates for an example session. The template color was randomly drawn on each block of trials, but with a reduced likelihood of choosing colors close to the previous template in order to uniformly sample the color wheel during each session. (**E**) Learning curves for monkeys B (top, 69 blocks) and monkey S (bottom, 102 blocks). The probability to choose the previous best target (e.g., pink) decreased after the template switch, while the probability to choose the current best target (i.e., green) increased. Solid lines and shaded areas represent the mean ± SEM probability of choice in monkey behavior. The dashed lines and shaded areas represent the model’s mean ± SEM probability of choice. Because there are 3 targets, chance level is 1/3. (**F**) Mean absolute angular distance between the estimated template (the color with the highest expected value according to the model) and the true template color decreased as monkeys progressed in the block (mean ± SEM across blocks, 69/102 blocks for monkey B/S, 30 bins using the normalized block length). (**G**) Examples of evolution of the expected value function. The brighter the color, the higher the expected value according to the model. Previous and current templates are indicated with dashed lines. The peak of the expected value function is indicated with a white marker on each trial. Before the template switch, the expected value function peaked around the previous template. After the switch, the estimated template either smoothly drifted toward the current template (*top*) or the expected value function reset to 0 (*bottom*, red line) causing the estimated template to abruptly move toward the current template. Resets were triggered by a large RPE, which typically occurred shortly after a block switch with a large change in the template color (Fig. S1D-E, see methods for details).

During each session, the template color changed repeatedly (Fig. 1B-D). Importantly, the monkeys were never explicitly instructed as to the identity of the template color or when it changed. Instead, they learned the template through trial and error, using reward as feedback to update their internal model of the template. Both monkeys quickly learned new templates: the proportion of trials in which the animal chose the best stimulus increased over the first dozen trials (Fig. 1E, solid green; performance was above chance after 15/7 trials for monkey B/S, p<0.05, Bonferroni corrected one-sided binomial test). As the animals learned the new template, they forgot the previous one, indicated by a decrease in the proportion of trials in which the animal chose the stimulus closest to the previous template (Fig. 1E, solid pink).

On average, the template was constant for a block of ∼100 trials (96 ± 53 / 88 ± 50 attempted trials, mean ± standard deviation, for monkey B/S, full range was between 30 and 306 trials). The change in template was not cued and occurred when the animal reliably chose the best available color (performance greater than 80/85% of 30 consecutive trials for monkey B/S, respectively). This allowed the monkeys to learn between 5 and 16 different templates per session (see Fig. 1D for example succession of templates; monkey B: 8 sessions, 8.6 ± 2.7 templates per session, monkey S: 9 sessions, 11.3 ± 3.4 templates per session). Note that, because the animals learned several templates, a particular stimulus feature that was the target on one trial could be a distractor on another, such that the animal had to use attention to search the array for the current best stimulus on each trial (Fig. 1B-C).

To model the monkey’s ability to learn an attentional template, we extended the standard Q-learning reinforcement learning model to approximate a continuous circular value function^37^ (see Fig. S1A for an example and methods for details). In this model, the attentional template was represented by the expected value of colors across the color wheel. On each trial, the model selected the stimulus with the largest expected reward. Then, after receiving reward feedback, the model updated its estimated value function such that colors with a positive reward prediction error (+RPE; i.e., they received more reward than expected) incrementally increased their value, while colors with a negative RPE (-RPE; i.e., received less reward) incrementally decreased their value (Fig. S1B shows an example update). In this way, the model captured the monkey’s ability to represent an attentional template, apply it to the stimulus array, and update the template based on feedback (Fig. 1E-F).

The model fit the monkey’s behavior well (Fig. 1E, dashed; model’s predicted likelihood of the chosen stimulus was 0.5826 ± 0.2635 / 0.5858 ± 0.2686 for monkey B/S). During learning, the probability that the animal and model chose the best option was highly correlated (p<0.001, r(80) = 0.8908 / 0.8697 for monkey B/S). Similarly, the model captured the animal’s forgetting the previous template (p<0.001, r(80) = 0.8687 / 0.8403 for monkey B/S). Finally, the model captured the relationship between the magnitude of the change of template color and the performance in the first 35 trials: the smaller the change, the better the performance (Fig. S1C). In fact, the model suggested monkeys used two strategies for learning a new attentional template after a change (Fig. S1D-E). Following a small change in the template color (< π/2), the animal incrementally updated their template (example shown in Fig. 1G, top). However, following a large change in template color (> π/2), the model predicted the animals ‘reset’ their attentional template (to uniform across colors) before re-learning a new template (Fig. 1G, bottom, see methods for details).

Overall, the animal’s behavior and the behavioral model suggest monkeys incrementally learned an attentional template and then used this template to guide their search and their decision to select a stimulus. With this foundation, we next aimed to understand how attentional templates are represented in the brain, how they are updated, and how they are used to guide decisions.

### Attentional templates were distributed across parietal and prefrontal cortex

To understand how attentional templates are represented across the brain, we recorded from prefrontal cortex, including lateral prefrontal cortex (LPFC, 492 neurons) and the frontal eye fields (FEF, 231 neurons), and parietal cortex (lateral intraparietal cortex, LIP, 167 neurons; Fig. 2A). We leveraged the Q-learning behavioral model to estimate the animal’s latent representation of the template on each trial. .Neurons in all three regions varied their activity as a function of the estimated template, taken as the color with the maximum expected value (Fig. 2B, see Fig. S2A for example model variables). Across the population, a significant number of neurons encoded the color of the estimated template (Fig. S2B; 30.54/40.26/30.08% in LIP/FEF/LPFC, all p≤0.002, permutation test, see methods for details). Given the structure of the task, this is the simplest way to represent the attentional template – one merely selects the color that is closest to the template. It is also consistent with the concept of a single attentional template^1–5^.

**Fig. 2.**
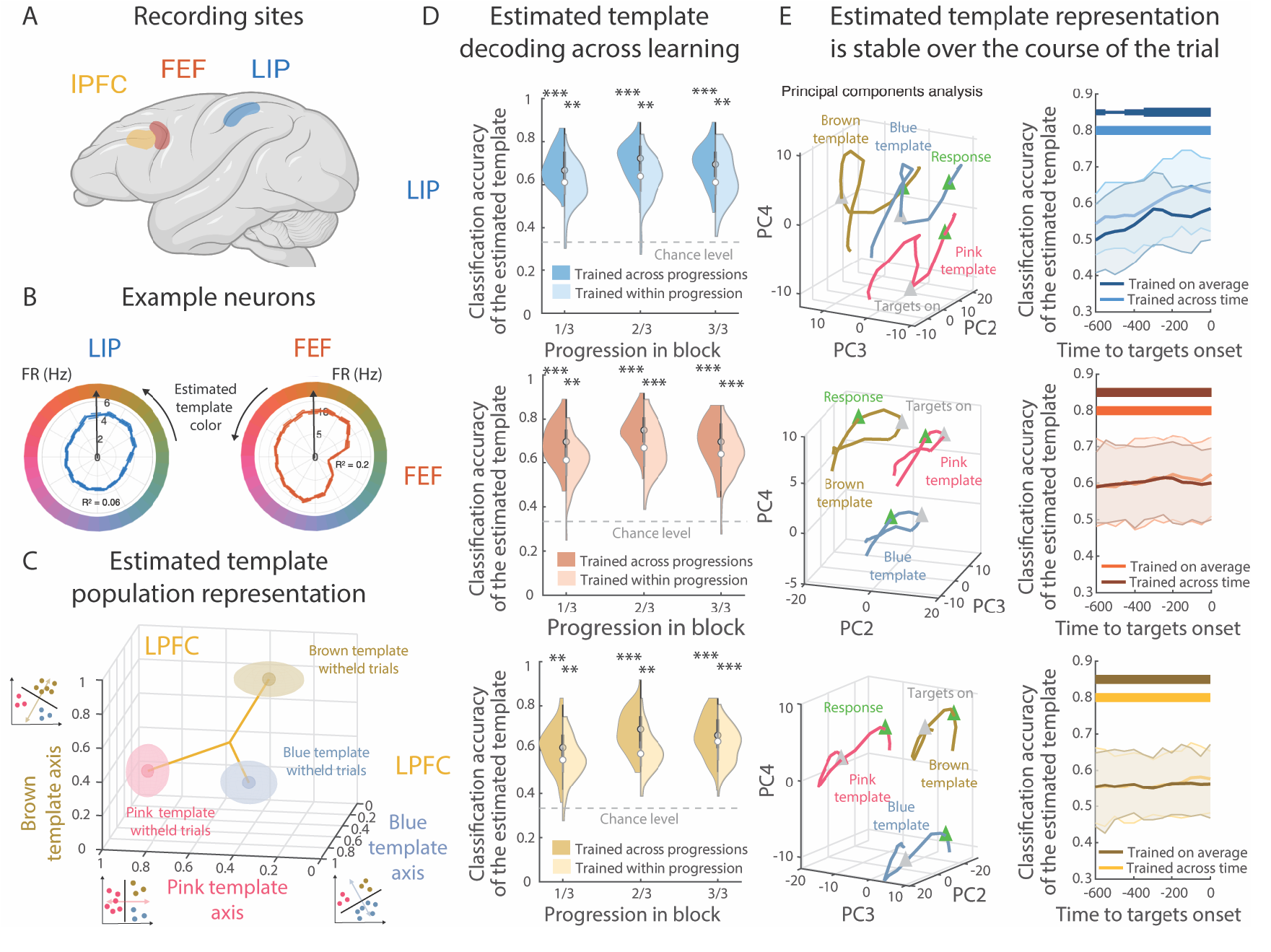
Distributed representation of the expected attentional template in the parietal and frontal cortex. (**A**) Recording sites. (**B**) Example neurons’ firing rate (mean ± SEM (dashed lines) across trials in a -600 to 300ms window around the onset of the targets) per bin (a sector of the wheel) of estimated template (bin size = π/6, smoothing of three bins) in LIP (blue) and FEF (red). (**C**) Neuron responses in LPFC (same time window as B), projected onto the vector normal to the hyperplane for the three classifiers trained to discriminate pink, brown, and blue templates (stratified to 100 neurons, trained on 61 trials per progression level (three levels) and estimated template color bin (three bins)). Next to the axes is a schematic representation of the pseudo-population decoding analysis. We split the estimated template in three color bins (brown, pink and blue) and trained a classifier for each color. Ellipses represent the mean (central dot) and the SEM (shaded) of the projection across the 100 bootstraps. The color of the ellipse corresponds to the color of the estimated template on withheld trials. (**D**) Classification accuracy of the estimated template (classifiers as in C) for each third of the block computed on withheld trials. Classifiers were either trained across progression levels in the blocks (left half) or within each progression level (right half). Violin plot^38^: central white dot is the median, thick vertical grey bar represents the 25^th^ to 75^th^ quartile and area represents the kernel density estimate of the data. (**E**) *Left*. The vector of activity in the neural population for each bin of estimated template color was projected into the reduced dimensionality space (of the second, third, and fourth eigenvectors in decreasing order of explained variance). LIP: 146 neurons, FEF: 216 neurons, LPFC: 475 neurons. Color code corresponds to the color of the estimated template. The grey triangle represents the onset of the targets and the green triangle the approximate time of response (200ms after the onset of the targets. The projection along the 4^th^ eigenvector shows the separation of the three estimated templates color in a triangle shape with a linear effect of time. The projection in the first 3 eigenvectors leads to a more cyclic effect of time (Fig. S2C-D). *Right*. Classification accuracy of the estimated template in 300ms windows either trained on the average activity in the full window (same as D and E) or trained for each window separately (100 bootstraps, mean and 95% confidence interval). For all panels, *p≤0.05, **p≤0.01 and bar thickness indicates significance level: p≤0.01 uncorrected, p≤0.05 Bonferroni corrected across time (13 time-points) and p≤0.01 Bonferroni corrected.

In addition to the estimated template, a subset of neurons represented the expected value of individual colors (Fig. S2B; 29.95/16.02/24.59% of neurons in LIP/FEF/LPFC all p≤0.002; see methods for details on model comparison). Such a continuous representation of expected value could allow one to estimate the expected value of any individual stimulus and, thus, calculate the RPE. Finally, a small proportion of neurons encoded the mean value of the trial or the chosen color (Fig. S2B). Here, we focus our analyses on the representation of the estimated template (i.e., the peak of the expected value function) as it was the most strongly represented in all three regions (Fig. S2B) and because it was the most compact representation of the template.

First, we were interested in understanding how the representation of the estimated template evolved during learning. While all three regions represent the estimated template, one might expect differences during learning (e.g., early representation in prefrontal cortex and later representation in parietal cortex). To track the evolution of template information, we trained a classifier to decode the estimated template from the neural population (see methods for details). In brief, we trained a set of three linear classifiers to discriminate the vector of neural response when the estimated template was in each of three bins (i.e., each classifier discriminated one third of the color wheel from the other two thirds, Fig. 2C, insets). All three classifiers decoded the estimated template on the majority of withheld trials in all three regions (e.g., Fig. 2C for LPFC). The output of these classifiers was then fed into a shallow feed-forward neural network that provided a single classification (see methods for details). This approach found all three regions consistently represented the template throughout learning (Fig. 2D, p<0.01 for all three regions, both across and within progression levels, see methods for details).

The estimated template was also stably represented during the trial. Plotting the neural activity in a low-dimensional space showed temporal dynamics within the trial were largely independent from the estimated template representation in LPFC and FEF (Figs. 2E, left column, and S2D). Consistent with this, classifiers trained to decode the estimated template during one time point of the trial (e.g., during the fixation period) could decode estimated template representations later in the trial (e.g., during response) in FEF and LPFC (Fig. S2E). In LIP, the coding of the estimated template was more dynamic, and the decoding accuracy increased specifically before and around the onset of the targets (Figs. 2D, right column, and S2E). However, in all three regions, classifiers trained on neural activity averaged over the trial’s entire time period could decode the estimated template at any individual time period (Fig. 2E, right column, all p<0.01; paired difference between training across and within each time period were all p>0.15). We did not find any differences in the timing or magnitude of the encoding of the estimated template across regions (Fig. 2D-E).

Together, our results suggest attentional templates were stably represented across frontal and parietal cortex, even as the animal learned (and relearned) new attentional templates. The stability of the representation may be important for allowing the template to guide attention on every trial. Next, we were interested in understanding the geometrical structure of the template representation.

### Neural representation of attentional templates was structured

As with many sensory features, neurons in sensory cortex that are selective to color tend to have smooth tuning curves (i.e., they respond similarly to similar colors)^39^. At the level of the neural population, this smoothness creates a structural geometry, such that perceptually similar colors are represented in similar ways in the brain^40,41^. However, it is unknown whether control representations have similar structure. In particular, we were interested in testing whether the geometry of the featural attention representation follows the shape of perception.

Structured template representations could facilitate generalization – by representing similar templates in similar ways, the brain could interpolate to new templates^42^. However, this would require a mechanism to constrain learning to a specific subspace, and therefore, could lead to interference when learning a new template. An alternative hypothesis is that cognitive control representations are high dimensional^43^ and so a unique neural representation is learned for each attentional template (regardless of perceptual similarity). Unique representations may be easier to learn, as template representations are not constrained, and high-dimensional representations would avoid interference between templates^44^.

To test whether the representation of attentional templates was structured, we revised the template classifier used above so that the neural network predicted the angle of the estimated template along the color wheel (Fig. 3A, see methods for details). Given the distributed nature of the template representation, we trained session-specific classifiers on the activity of simultaneously recorded neurons from all three regions (this also increased the classifier’s statistical power). We then applied the classifier to the first 35 trials of a new template color (all withheld from training) to test two predictions of a structured representation (Fig. 3B).

**Fig. 3.**
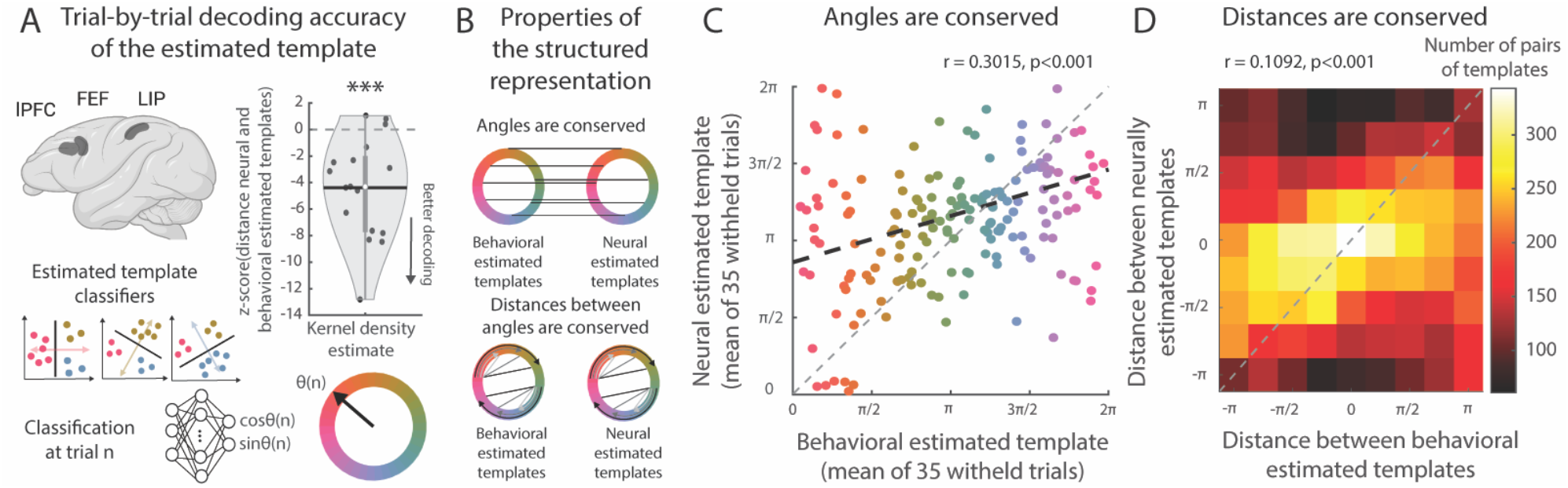
Neural attentional templates are structured. (**A**) To decode the trial-by-trial estimated template, we combined the neural activity of the three regions of interest. We used three classifiers trained to classify each color bin. The posterior probability estimated by all three classifiers were passed as input into a feedforward, fully-connected, neural network trained to predict the cosine and sine of the estimated template. Thus, on each trial n, we could estimate the color *θ*(*n*) of the neural estimated template. Violin plot: mean z-scored circular distance between the neural and behavioral estimated template on withheld trials for each session. Black dots are individual session, central white dot is the median, horizontal black bar is the mean, thick vertical grey bar represents the 25^th^ to 75^th^ quartile and area represents the kernel density estimate of the data. (**B**) Schematic of properties of a structured representation. (**C**) Mean neural and behavioral estimated template across blocks, color is that of the mean behavioral estimated template (r = 0.3015, p<0.001, 171 templates, permutation test on the circular distance). Thick dashed line indicates the circular correlation. (**D**) Histogram of the circular distances between mean neural and behavioral estimated templates across blocks (r = 0.1092, p<0.001, 14,535 pairs, permutation test on the circular distance).

First, if the neural representation of the template is structured, then it should match the behavioral template (which was structured by construct). Indeed, the neurally estimated template was correlated with the estimate of the behavioral model (r(5,838) = 0.1788, p<0.001, circular correlation) and the distance between the neural and behavioral template was significantly less than expected by chance (average absolute circular distance = 1.2662, p<0.001, permutation test). Note, all analyses were done on withheld trials. This effect was consistent across sessions (Fig. 3A; t(16)=-4.7694, p<0.001, one-sided t-test across sessions). Furthermore, as seen in Figure 3C, the neurally-estimated color was correlated with the behaviorally-estimated color across the spectrum (r(171) = 0.3015, p<0.001, circular correlation; absolute circular distance = 1.1419, p<0.001, permutation test). Interestingly, the decoding error of the neural model increased when the animal’s behavior suggested they were uncertain about the attentional template (measured as the entropy of the expected value function, Fig. S3A-B).

Second, a structured neural representation should capture the semantic relationship between templates – templates close in perceptual space should also be close in neural space (Fig. 3B, lower). Consistent with this, we found the distance between pairs of mean neural estimated templates could predict the distance between behavioral estimated templates (Fig. 3D, and Fig. S3C for all pairs; r(14,535) = 0.1092, p<0.001, circular correlation; absolute circular distance = 1.4267, p<0.001, permutation test).

Overall, our results suggest frontal and parietal cortex represent the monkeys’ internal model of the current attentional template in a structured fashion, with perceptually related templates represented in similar ways. Next, we were interested in understanding how the neural representation of the attentional template was learned.

### Attentional templates in frontal and parietal cortex were learned through incremental updates

Several mechanisms are thought to support learning of neural representations. Unsupervised learning is thought to shape representations in early visual cortex^45^. Predictive (semi-supervised) learning can capture representations of language^46^. Supervised, reward-driven, reinforcement learning is thought to underlie associative learning^47,48^. However, relatively little is known about how attentional control representations are learned.

The animal’s behavior suggests templates may be learned using reinforcement learning in a continuous space (Fig. 1E). If true, then this makes several predictions about how the attentional template should change in response to feedback. First, when the animal receives a reward that is higher than expected (+RPE), then the template representation should shift to be more similar to the chosen color. This effect was observed in the animal’s overall behavior (t(2,945)=15.0389, p<0.001, one-sided t-test, see methods for details) and was consistently observed across individual sessions (Fig. 4A; mean behavioral update was toward the chosen color, one-sided t-test: t(16) = 13.4583, p<0.001, one-sided t-test). A similar effect was seen in the neural template. Following a +RPE, the neural template was updated such that it shifted towards the chosen color (t(2,945)=4.2168, p<0.001, one-sided t-test). Again, this effect was consistent across sessions (t(16) = 3.7321, p<0.001, one-sided t-test). Furthermore, the behavioral update predicted the neural update (p=0.014, by permutation test on absolute circular distance), an effect that was consistent across sessions (Fig. 4B, t(16)=-3.3410, p=0.0021, one-sided t-test).

**Fig. 4.**
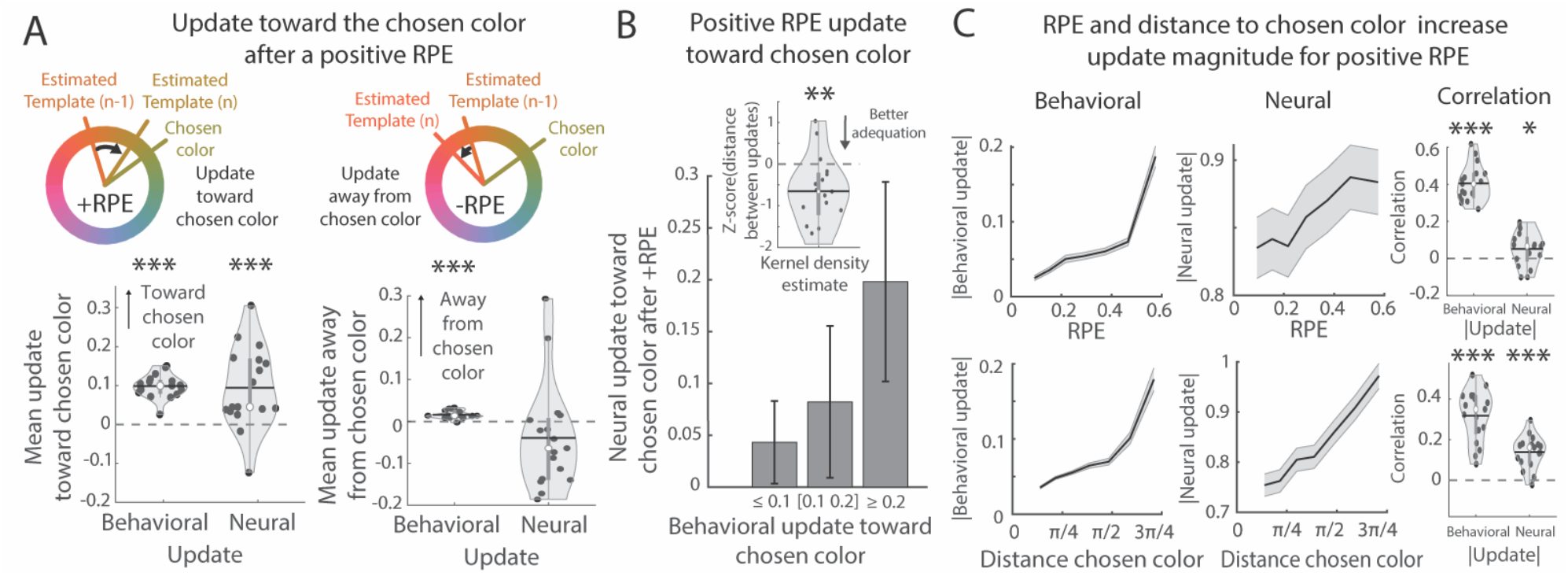
Neural attentional templates are incrementally updated in the structured space. (**A**) *Top-left*. Example model prediction for a positive reward prediction error (+RPE). The estimated template moves toward the chosen color. *Bottom-left*. Mean behavioral and neural are updated towards the chosen color across sessions (see methods, all on withheld trials). *Right*. Same as left, but for negative RPEs (-RPE) where the estimated template moves away from the chosen color. (**B**) Mean neural update toward the chosen color (± SEM) after +RPE as a function of the size of the behavioral update (865, 268 and 170 trials in each bin). Inset shows violin plot of mean z-scored distance between the neural and the behavioral update toward the chosen color across validation trials for each session. (**C**) *Top*. Mean neural (*left*) and behavioral (*middle*) update magnitude (± SEM) for each bin of RPE magnitude (10 bins, smoothing of 4 bins) across all validation trials following +RPE. Violin plot (*right*): mean Pearson correlation between the update magnitude and the RPE magnitude across validation trials for each session. *Bottom*. Same as *top* but for absolute distance between the estimated template before the update (at n-1) and the chosen color. For all panels, * p≤0.05, ** p≤0.01, *** p≤0.001.

A second prediction is that, if the neural representation of the template followed a reward-driven update rule, then the change in the representation should be larger when a) the animal received a greater magnitude +RPE or b) the distance between the previous template and the (rewarded) chosen color was greater. As predicted, the behavioral update depended on the +RPE magnitude (Fig. 4C, left; r(2,945)=0.3676, p<0.001, one-sided Pearson correlation), an effect that was consistent across sessions (Fig. 3G, right; t(16)=17.0713, p<0.001, one-sided t-test on correlation coefficients). Although weaker, a similar effect was seen for neural updates: the overall effect was trending (Fig. 4C, middle; r(2945)=0.0243, p=0.0939, one-sided Pearson correlation) but the effect was consistent on individual sessions (Fig. 4C, right; t(16)=2.2864, p=0.0181, one-sided t-test). Second, as predicted, the update magnitude was related to the distance between the template and chosen color for both behavioral and neural updates (Fig. 4C, middle; behavioral: r(2,945)=0.2941, p<0.001; neural: r(2,945)=0.1422, p<0.001, one-sided Pearson correlation). Again, these effects were consistent across sessions (Fig. 4C, right; behavioral: t(16)=10.3213, p<0.001; neural: t(16)=7.1118, p<0.001, one-sided t-test). Consistent with the update in template being influenced by both +RPEs and the distance between template and choice color, the greatest changes in template representations were observed after the template changed, when learning was greatest (Fig. S4A).

For negative RPEs (-RPE), the behavioral model predicted a small update that would shift the template away from the chosen color (Fig. 4A; t(2,669)=14.3303, p<0.001; t(16)=7.1538, p<0.001 across sessions, one-sided t-test; this effect was ∼10x smaller than +RPE). Interestingly, we did not observe this in the neural data (t(2,669)=-2.1542, p=0.9843; t(16)=-1.2796, p=0.8905 across sessions, one-sided t-test). And, unlike +RPEs, the magnitude of the update in the neural template was not related to -RPE magnitude nor distance between template and chosen color (Fig. S4B). We do not think that the lack of update following a -RPE was because the learning rate was different: models that included different learning rates for +RPE and -RPE did not fit the behavior better than the simpler model with one learning rate (Monkey B: ΔBIC = 467, Monkey S: ΔBIC = 406, see methods for details). Instead, the lack of a clear update in the neural template following a -RPE could reflect low signal to noise ratio (due to the reduce influence of value on choices) or a different mechanism for responding to -RPEs (such as inducing exploration)^49^.

Altogether, these results suggest the representation of the estimated template in frontoparietal cortex was learned through a series of incremental updates following each trial. By repeatedly shifting the template towards stimuli with +RPEs, the template could evolve towards the optimal template color for that block of trials.

### Prefrontal and parietal cortex represented the value of individual stimuli in a way that generalized across templates

In the current task, the attentional template must be combined with sensory information in order for the animal to decide which stimulus to select with a saccade (i.e., overt attention). There are two hypothetical models for how templates could guide decisions. First, the attentional template could change the decision process. In this ‘decision modulation’ model, the decision boundary changes for each template, rotating to optimally discriminate stimuli close/far from the template boundary (i.e., those with high/low value; Fig. 5A, left). The decision boundary could be learned incrementally, similar to the template, or chosen from a set of pre-determined decision boundaries^50^. Alternatively, in a ‘generalized value’ model, the attentional template acts on sensory representations in a way that transforms them into generic ‘value’ signals (Fig. 5A, right). Although this would require re-mapping stimulus responses for every template (e.g., both ‘pink’ when the template is ‘red’ and ‘teal’ when it’s ‘blue’ are high value), it would have the advantage of only needing one, stable, decision boundary^51–53^. Several lines of evidence support the second hypothesis.

**Fig. 5.**
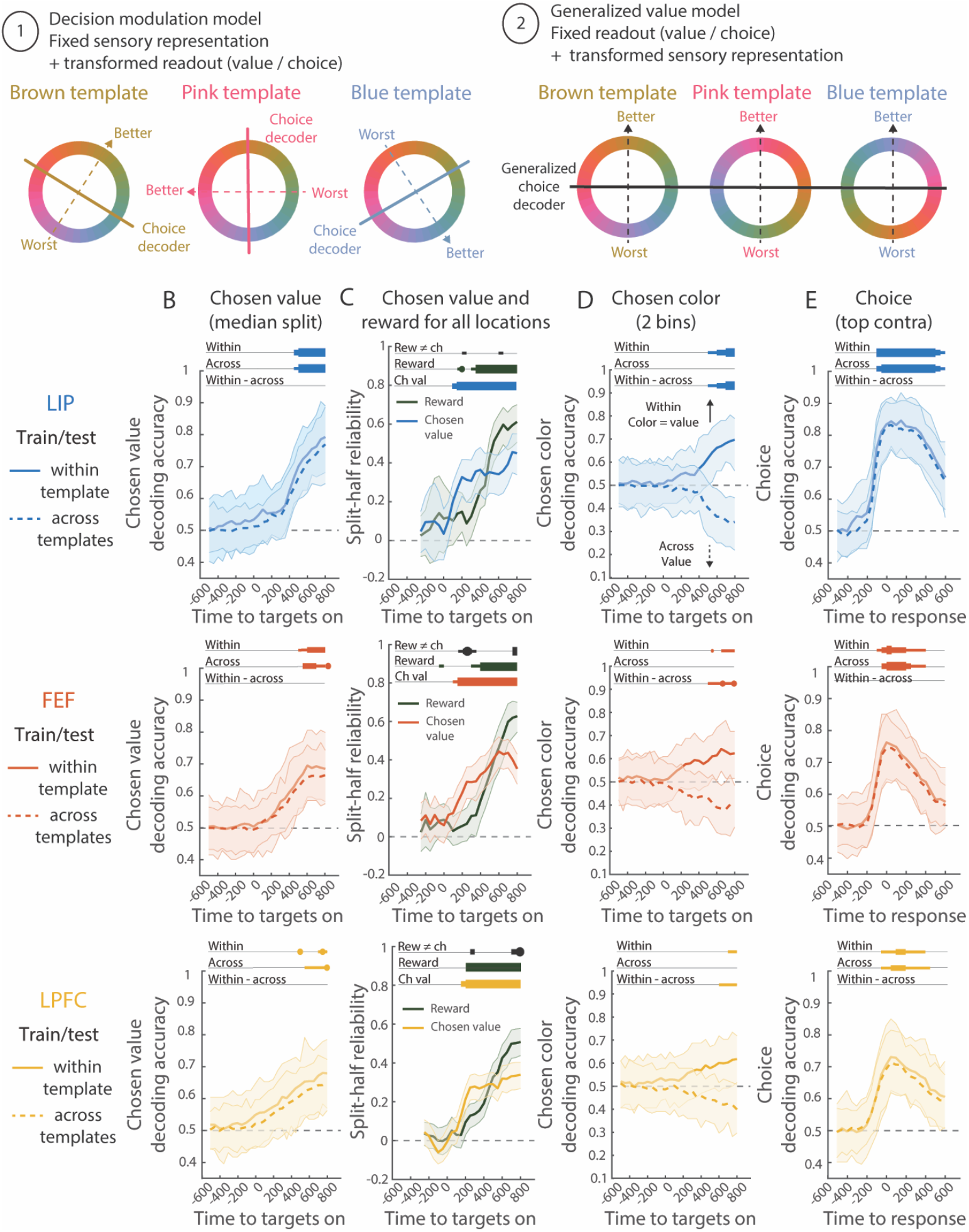
Sensory-agnostic stimulus re-mapping for a generalized decision-making process across templates. (**A**) Hypotheses for how the attentional template is used to make a choice. 1) Decision modulation model: the network has a fixed representation of the sensory perception; the readout (choice and value) rotates based on the attentional template to determine which color matches the template; 2) Generalized value model: the network has a fixed readout (choice and value) and transforms the sensory perception based on the template to indicate which colors are desirable. (**B**) Classification accuracy (with 95% confidence interval) of the chosen value over time (median split within each session). Computed on withheld trials that were in the same estimated template color bin as the training trials (within, solid line) or in a different estimated template color bin (across, dashed line; 110 neurons, training on 128 trials per estimated template bin, three bins). Chancel level was 1/2. Bar thickness indicates significance level with increasing line thickness: p≤0.01, p≤0.05 Bonferroni-corrected across time (27 time points) and p≤0.01 Bonferroni-corrected. (**C**) Time course of the mean split-half reliability (with 95% confidence interval) of the chosen value and reward regressors computed across all locations (see methods for details). Bars indicate the significance of the split-half reliability and the difference in reliability strengths between the two regressors (black): p≤0.01, p≤0.05 Bonferroni-corrected across time (22 time points) and p≤0.01 Bonferroni-corrected. (**D**) Same as B, but for classifiers trained to decode the chosen color (balanced for the estimated template, 2 bins; 133 neurons, trained on 160 trials per estimated template bin). Chance level was 1/2. Significance level threshold was lowered: bar thickness indicates significance level: p≤0.05, p≤0.01 and p≤0.05 Bonferroni corrected across time (27 time points). (**E**) Same as B, but for decoding choice at the best contralateral location (90 neurons, training on 120 trials per estimated template bin, three bins). Chancel level was 1/2. Bar thickness indicates significance level: p≤0.01, p≤0.05 Bonferroni corrected and p≤0.01 Bonferroni corrected. Bonferroni-correction was across time (23 time points) and locations (4 locations).

First, the value of the stimulus chosen by the animal was represented in a stable manner that generalized across templates. A classifier trained to discriminate between chosen stimuli with high and low expected values performed significantly above chance in all three regions (Fig. S5A, note that this controls for movements, see methods for details). Critically, the same classifier also performed above chance on blocks of trials with a different attentional template (Fig. 5B). In fact, the classifier performed equally well on the template it was trained on and when tested on different templates (difference between templates was not significantly different from zero, p≥0.24 uncorrected for all three regions, permutation test, see methods for details).

Similar results were found when fitting generalized linear models (GLMs) to estimate how strongly neurons represented the value of the chosen stimulus, while controlling for potentially confounding variables such as reward (see methods for details). The value of the chosen stimulus was represented significantly in all three regions (Fig. 5C; measured with the split-half reliability of the model^54^, see methods for details). This representation occurred immediately after the presentation of the stimuli, preceding the representation of the reward (Fig. 5C, p=0.001 in FEF and p=0.044 in LPFC, with a weaker effect in LIP, p≥0.065, all Bonferroni-corrected). These results suggest neurons represent the value of the chosen stimulus, consistent with the generalized value model.

In addition, the colors of the stimuli were not encoded in any of our recorded regions. Decoders trained on the responses of the neural population failed to decode the color of a stimulus at any location in the visual array (Fig. S5B) and failed to decode the color of the selected stimulus (Fig. S5C, controlling for the estimated template color). This is consistent with our observation that the color of the chosen stimulus was represented in very few or no individual neurons (Fig. S2B-C). In line with a remapping of the sensory representation in the frame of the estimated template, we found a weak representation of the chosen color for a given template (Fig. 5D; p=0.03 Bonferroni-corrected in LIP, and trending in FEF and LPFC, p=0.02/p=0.03 uncorrected, see methods for details) and these decoders did not generalize to trials with a different template (Fig. 5D, p≥0.42 uncorrected for all three regions; decoding was worse than within the same template in p=0.018 Bonferroni-corrected in LIP and trending in FEF and LPFC, p=0.007/p=0.016 uncorrected). If anything, when comparing across templates, color decoding was below chance, as would be expected if the chosen color re-mapped when the template changed (as predicted by the generalized value model).

Finally, we tested whether a classifier trained to discriminate between chosen and unchosen stimuli generalized across attentional templates. Such ‘decision’ decoders found strong encoding of choice in all three regions (Fig. S5D, see methods for details). Consistent with a static decision boundary, classifiers trained on trials from one attentional template generalized to other templates (Fig. 5E and Fig. S5E for all locations). In fact, classifiers performed equally well on withheld trials from the same template or another templates (Fig. 5E; difference between within and across templates was not significantly different from zero, p≥0.2232 uncorrected for all three regions). It is important to note that these decision classifiers likely integrate both the decision process and the execution of the movement. While movements are expected to generalize across templates, the fact that we observed generalization at all time periods, including at the earliest moments before the saccade, suggests the decision-making process also generalizes across templates (as predicted by the generalized value hypothesis).

Altogether, these results are consistent with the hypothesis that the attentional template transforms sensory inputs into a generalized value representation in prefrontal and parietal cortex. The result is a ‘priority map’ of the value of the stimulus at each location^55,56^. By generalizing across templates, such value representations could allow the same neural circuitry to be used to decide which stimulus to select, regardless of the attentional template. This avoids the need to learn bespoke decision circuitry for each template and allows the brain to flexibly control attention in many different tasks^51–53^.

### Value representations transform from location-specific to global

Finally, we were interested in understanding how these value representations support decisions and learning. When making a decision about where to look, the animal must compare the value of stimuli across locations in order to identify and select the stimulus with the greatest expected value. This computation requires a ‘local’ representation of the value of the stimulus at each location. However, learning the attentional template should be agnostic as to the location of the stimulus and, thus, requires a ‘global’ representation of value. To understand how value was represented, we fit a generalized linear model (GLM) to quantify the encoding of the value of chosen and unchosen stimuli both ‘locally’ (at each location) and ‘globally’ (across locations; all comparisons used split-half reliability^54^, see methods for details). Our analyses revealed value representations were dynamic, initially local to each location (to support decision-making) before evolving into a global representation (to support learning).

Immediately after stimulus presentation, the value of a stimulus was represented in a location-specific manner. In particular, value representations were initially restricted to the contralateral hemifield (consistent with hemispheric lateralization during attention^7,9,57^). LPFC represented the value of stimuli in the contralateral hemifield more strongly than ipsilateral stimuli, regardless of whether it was chosen or unchosen (Fig. 6A-B; although this was weaker in FEF or LIP, Fig. S6A-B; permutation test, see methods for details). In fact, there was almost no encoding of the value of unchosen ipsilateral stimuli at any point in the trial (Fig. 6B and S6B, see Fig. S6C-D for comparison between areas). Further reflecting a localized representation, the neural representation of the value of chosen stimuli were more similar for locations that were closer together in physical space (Figs. 6C and S6E-G, measured as the correlation in beta-weights, permutation test).

**Fig. 6.**
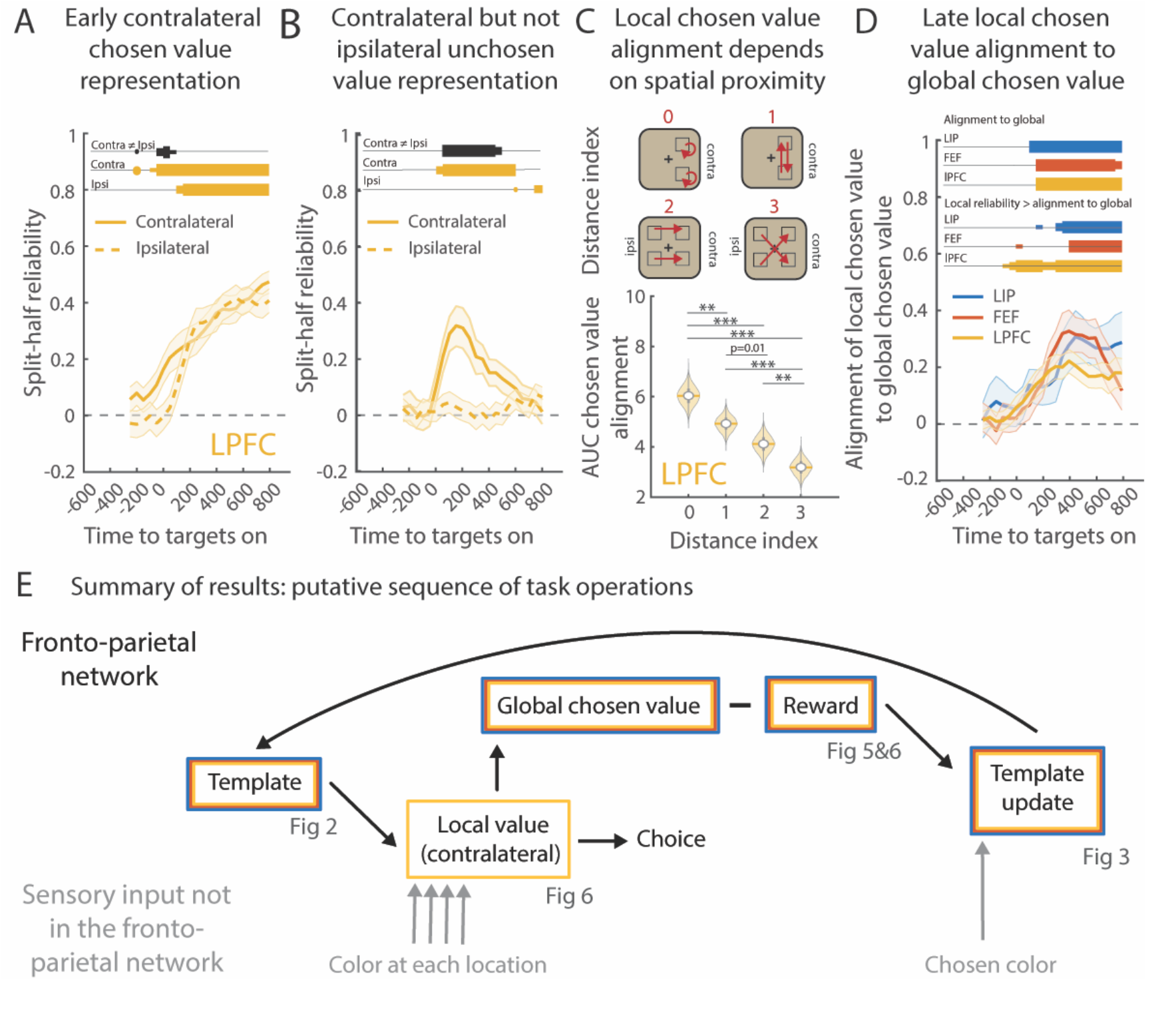
Value representations transformed from location-specific to global over time. (**A**) Time course of the mean split-half reliability (with 95% confidence interval) of the chosen value regressors for the two contralateral locations (contra) and the two ipsilateral locations (ipsi, dashed line) for LPFC (see Fig. S5A for LIP and FEF). Bars indicate the significance of the split-half reliability (yellow) and the difference in reliability strengths between the two regressors (black). (**B**) Same as A, but for the unchosen value regressors (see Fig. S5B for LIP and FEF). (**C**) *Top*. Distance index: the source of the arrow indicates the location where the chosen value representation was estimated in one half of the session and the head of the arrow where it was applied (in the other half of the session). *Bottom*. Violin plot: mean area under the curve of the representation reliability (distance index = 0) or alignment across locations across time (22 time points, Fig. S5E) across bootstrap for LPFC. Central white dot is the median, horizontal bar is the mean, thick vertical grey bar represents the 25^th^ to 75^th^ quartile and area represents the kernel density estimate of the data. Statistical significance was estimated using a paired one-sided z-test (assuming that smaller distances would be more similar) using the reliability or the correlation disattenuation against 0. * p≤0.05, ** p≤0.01, *** p≤0.001 Bonferroni-corrected (6 pairs). See Fig S6G or LIP and FEF. (**D**) Same as A, but for the correlation between the contralateral chosen value representation (shown in A and Fig. S5A) and the global chosen value representation (shown in Fig. 5C) for LIP, FEF and LPFC. Upper bars (Alignment to global) indicate the significance of the correlation disattenuation between the two vectors. Lower bars (Local reliability > alignment to global) indicate whether the correlation between the two vectors is significantly lower than the local chosen value reliability. (**E**) Summary of results and putative sequence of task operations. For all panels, bar thickness indicates significance level: p≤0.01, p≤0.05 Bonferroni-corrected across time (22 time points) and p≤0.01 Bonferroni-corrected.

If these initial, location-specific, representations of value provide the input to the decision-making process, then they should proceed the decision to choose a stimulus. This predicts the initial representations should be similar for both the chosen and unchosen stimuli. Consistent with this, the representation of the value of the chosen and unchosen stimuli were similar to one another in LPFC (measured as correlation of beta-weights, p=0.0341 Bonferroni-corrected in a 200ms window centered on 150ms after the onset of the targets, permutation test, see methods for details). This similarity was also seen in LIP and FEF, although it was weaker (p=0.094/0.0598, uncorrected).

After initially being represented in a local manner, LPFC began to represent the value of ipsilateral stimuli, but only when they were chosen (Fig. 56A-B). This suggests the value of the chosen stimulus was initially local but evolved into a ‘global’ representation that generalized across locations. Consistent with this, the local representation of value slightly proceeded the global representation (local: Fig. 6A, first significant in 200ms window centered on 50ms in LPFC; global: Fig. 5C, 250ms; all Bonferroni corrected). Furthermore, the representation of the value of the contralateral stimuli (i.e., the ‘local’ representation) changed over time, becoming increasingly aligned with the global representation (Fig. 6D, measured as correlation in GLM beta weights; significant at the time that the global representation emerges, 100/150/150ms after the onset of the targets in LIP/FEF/LPFC, permutation test, see methods for details). Finally, the representation of the chosen value at each location a) became aligned with the representation of the chosen value at other locations (Figs. 6C and S6E-G) and b) could significantly explain variance in the chosen value at other locations (Fig. S6F-H).

Altogether, Figures 5 and 6 show the attentional template transformed the representation of each stimulus into a local representation of the value at that location. This local representation is necessary for decision-making processes to determine where to move one’s eyes. Then, over time, the local representation transformed into a global, location-agnostic, representation (particularly in LPFC, Fig. 6). Such a global value representation could be used to calculate a reward prediction error, regardless of the location of the chosen stimulus. By abstracting away from a specific location, the global value could allow for the attentional template to be updated in a location-agnostic manner, allowing learning to generalize across stimulus locations.

## Discussion

Attention selects task-relevant sensory information for processing and action^8^. It is guided by an ‘attentional template’ that represents the set of stimulus features that are currently relevant^2–4^. Here, we aim to understand how the brain learns attentional templates and then uses them to influence sensory processing and drive decision-making processes. To this end, we trained two monkeys to perform a visual search task that required them to repeatedly learn new attentional templates (Fig. 1). Our results are summarized in Figure 6F, which provides a theoretical schematic of how attentional templates influence decision-making. In brief, we found the current attentional template (the estimated template in our model) was represented in a distributed fashion across prefrontal and parietal cortex (Fig. 2). Template representations were structured, with related templates (nearby in color wheel) represented in a similar manner (Fig. 3). During the visual search, the template acted on sensory representations such that the feature of the stimulus was converted into a ‘value’ representation (Fig. 5). Value representations generalized across templates, allowing the decision process of whether to select a stimulus to be the same across all templates (Fig. 5). To facilitate the decision of where to look, the value of each stimulus was represented in a topographic manner, with independent ‘local’ value representations at each stimulus location (Fig. 6). Once a stimulus was chosen, its value transformed from a location-specific representation to a ‘global’ representation that was agnostic as to the location of the stimulus (Fig. 6). We hypothesize this global representation could allow the brain to calculate a reward prediction error that generalized across the location of the chosen stimulus. Finally, we found this general reward prediction error was used to update the attentional template, such that attention shifted towards rewarded features (Fig. 4). Altogether, our results provide a broad understanding of how attentional templates are learned and how they are used to guide eye movements in the frontal and parietal cortex.

### Attentional templates are represented in a structured manner across frontal and parietal cortex

Consistent with previous work, we found attentional templates are represented in a distributed network that includes prefrontal and parietal cortex^28–35^. The distributed nature of these representations may reflect the fact that attention emerges from the dynamic interaction between prefrontal and parietal cortex. Alternatively, attention may be controlled from one region and then quickly broadcast to the other regions. Consistent with LPFC playing a leading role in attention, we found value representations were generally strongest in LPFC: the representation of the value of unchosen stimuli was stronger in LPFC than FEF (Fig. S6C, p=0.0041 Bonferroni-corrected, permutation test, see methods for details) and peaked earlier in LPFC and LIP than FEF (Fig. S6D, LIP: p=0.0217, LPFC: p=0.0355, permutation test, see methods for details). Future work is needed to causally test this hypothesis – recent work has shown inactivating contralateral LPFC impairs a monkey’s ability to choose a target based on feature-based attention^9,57^ but it is unknown if similar effects can be found when inactivating parietal regions.

Our results suggest the neural representation of attentional templates were structured, with perceptually similar colors represented by similar patterns of neural activity (Fig. 3). Note that this is not trivial – previous work has suggested high-dimensional or random representations may facilitate learning^43^ which would predict a unique attentional template on each block of trials. However, structured template representations may have several advantages that make them more useful for the brain. First, by matching the structure of sensory representations, moving through the space of control representations would have a smoothly varying effect on sensory representations. For example, a neuron encoding ‘red’ templates, that biases ‘red’ representations in sensory cortex, would have a graded response to ‘pink’ and ‘purple’. Second, structured representations could facilitate learning. When template representations vary smoothly, then updating the value of a single color should have a graded effect on nearby colors. This was reflected in the model fit to the animals’ behavior, which was well fit by a Q-learning with function approximation model that posits learning occurs over a small set of smooth functions that form a basis for representing a continuous space (Figs. 1 and S1). Third, representing attentional templates smoothly in neural space could allow for generalization to new templates. In particular, the continuous space of templates could allow the animal to interpolate a template that had never been experienced. For example, a new ‘rose’ template could be inferred to lie between ‘red’ and ‘pink’. One cost of a structured template is that, for a given feature, the brain may be limited to a single attentional template. Indeed, psychophysical studies suggest this may be true^58^ (but see^59^).

Future work is needed to understand how the structure in neural representations emerges. Structure could have been acquired during training, either because rewards were graded for nearby colors or as the animals learned the general ‘meta’ structure of the task^60–62^. Alternatively, structured organization could have been inherited from sensory cortex. Even when passed through random connections, neural representations maintain their local topography and so prefrontal and parietal regions may inherit structure from sensory cortex^44^. Consistent with such an innate structure, spatial representations are structured, even when animals are first learning a working memory task^63^.

### Attentional templates can be incrementally learned through reward-driven reinforcement learning

A long history of psychophysical studies has shown the importance of rewards in shaping attentional templates^17–23^. For example, previous work has shown reward modulates the representation of stimuli throughout the brain, including in prefrontal and parietal cortex^22^ and can lead to sharpening of tuning responses in visual cortex (similar to attention)^64^. Our work builds on this foundation to study the neural mechanisms by which reward can shape attentional templates. Both behavioral modeling and neural results suggest the brain uses reinforcement learning to update the internal model of the attentional template. Following a positive reward prediction error (+RPE), the attentional template was shifted towards the chosen color with a predicted magnitude (Fig. 4). These results are consistent with the large body of work showing reward learning can modulate the value of individual (discrete) representations^65^, including updating the belief about the current task^35,66,67^. Our work extends this to the continuous domain and provides insight into how new control representations could be learned.

Future work is needed to further detail the mechanisms of learning. Learning the value of individual stimuli is thought to rely on the representation of reward prediction error in dopaminergic neurons^68,69^. Given the strong dopaminergic innervation of prefrontal cortex, a similar mechanism may act to update control of internal and cognitive states^70^.

### Creating a ‘priority map’ for a generalized decision-making process

Despite being the task-relevant sensory feature, the color of stimuli was not represented in prefrontal or parietal cortex during our task (Figs. S2B-C, 5D and S5B-C). This is not because these regions are not sensitive to color: stimulus color information has previously been found in all three regions^41,71–74^. Instead, we found that featural attention re-mapped the sensory information into a ‘value’ representation that generalized across templates. In this way, prefrontal and parietal cortex represented a ‘priority map’ of the visual scene.

Priority maps represent the priority of each stimulus in the visual field^4,55,56,75^. This can be used to guide attention and decisions – in our task, the monkey simply selects the stimulus with the highest priority. These maps often reflect bottom-up, external characteristics of stimuli (e.g., size, brightness, etc.). Indeed, previous work has found priority maps of visually salient stimuli are represented in both parietal and prefrontal cortex^55,56^. However, as is the case in our task, priority maps can also integrate top-down, goal-directed, attention to specific stimulus features^4^. Our results show how attention integrates the current attentional template with sensory drive to form a map of the value of stimuli in the contralateral hemifield of the visual array. This map is then used to guide visual search and make a decision as to where to saccade in the visual scene (a form of overt attention).

Future work is needed to understand the mechanisms that generate a map of stimulus value. Previous work provides some possible mechanisms. Attention is known to increase the response of neurons that represent stimuli matching the attentional template^55,56^. Therefore, by summing the activity of the neural population representing each location in space, one could estimate how closely the stimulus at each location matched the attentional template (i.e., that location’s value). This would result in a priority map. Winner-take-all dynamics between locations on the priority map can then select the location with the greatest priority, driving attention (either overt or covert) to that location^55^.

### A common framework for attention, reward learning and value-based decision-making

As noted above, attention, reward-learning and value-based decision-making are thought to be closely interwoven cognitive operations. Our task requires attention, as the animals must use the template to perform a covert visual search for a matching stimulus while ignoring distractors (both hallmarks of attention). However, the task also requires value-based decision-making, as the animals must use reward outcomes to update their internal representation of the attentional template.

Although these concepts have been largely studied independently, our results show how integrating concepts from both domains can deepen our understanding of attention and decision-making. In our task, the mechanisms supporting learning attentional templates parallels the reinforcement learning mechanisms observed in simpler associative learning. Furthermore, our results show how value-based representations can allow the brain to generalize decision-making across a variety of attentional templates. These results suggest attention and value-based decision-making may be overlapping, with a shared set of neural mechanisms.

### Beyond attention, implication for learning new tasks

Attention is one form of cognitive control, the control of sensory processing. Our results may extend to other forms of cognitive control. For example, we found attentional templates are structured, such that semantically related tasks are represented in similar ways in the neural population. However, perception is not alone in being structured; previous work has shown memories, spatial layouts, cognitive maps, and conceptual knowledge are also represented in a structured manner^41,76–80^. Given this, cognitive control may be structured in other domains. This suggests there may be one or more continuous ‘task spaces’ in which smoothly changing the activity in the neural population may allow the brain to navigate through semantically related cognitive control states. As noted above, this could allow one to generalize to new situations by interpolating within task space.

In addition, our results suggest cognitive control can be learned through reinforcement learning, providing neural evidence for incremental learning of tasks^62^. Such a mechanism could learn cognitive control representations *de novo*, without any relationship to other tasks. However, if task representations exist within a continuous task space, then incremental learning could move cognitive control states through this space in order to flexibly adapt to a changing environment. This could be combined with other mechanisms for learning cognitive control. For example, when in a new situation, one might use contextual cues to recall a generally appropriate task representation from previous, similar, experiences (e.g., searching for a particular color car when looking for a cab). Incremental learning could then optimize the task representation for the current situation.

## Acknowledgements

The authors thank Pavlos Kollias for help building the behavioral task control system; Neeraja Rajagopalan, Matthew Panichello and Sina Tafazoli for help during recordings; Nathaniel Daw, Flora Bouchacourt and Harrison Ritz for their insights into the behavioral and neural analyses; and Alex Libby, Junchol Park, Harrison Ritz, Qinpu He, Sina Tafazoli, Motoaki Uchimura and Adel Ardalan for their feedback during the writing of this manuscript. This work was supported by NSF CAREER BCS-2143391 to TJB and a post-doctoral project grant by the Phillips’ Foundation to CIJ.

## Author Contributions

Conceptualization, TJB, RBE, CIJ; Investigation, CIJ, BM, NTM, and TJB; Formal Analysis and Visualization: CIJ.; Writing – Original Draft, CIJ and TJB; Writing – Review & Editing, CIJ and TJB; Funding Acquisition and Supervision: TJB.

## Methods

### Resources availability

The lead contact for this study is Timothy J. Buschman.

Both the code and neural data supporting each figure are publicly available online at https://github.com/caroline-jahn/LAT_062923.

This study did not generate new unique reagents.

### Key resources table

| RESOURCE | SOURCE | IDENTIFIER |
| --- | --- | --- |
| MATLAB | Mathworks | RRID:SCR_001622 |
| VBMC toolbox | Acerbi, 2018<sup>81</sup> | https://github.com/lacerbi/vbmc |
| Plexon Offline sorter | Plexon | https://plexon.com/products/offline-sorter/ |
| circular statistics | Mathworks | RRID:SCR_016651 |
| Violin plot | Bechtold et al., 2021<sup>38</sup> | https://github.com/bastibe/Violinplot-Matlab |

### Ethics

All experimental procedures were approved by the Princeton University Institutional Animal Care and Use Committee and were in accordance with the policies and procedures of the National Institutes of Health.

### Task and behavior

Two adult male rhesus macaques (monkey B, 13 kg, and monkey S, 9 kg) performed the experiment. Stimuli were presented on a Dell U2413 LCD monitor positioned at a viewing distance of 58 cm. The monitor was calibrated using an X-Rite i1Display Pro colorimeter to ensure accurate color rendering. The task consisted in choosing one of three colored targets on a screen. These three colors were randomly selected from 100 evenly spaced points along an isoluminant circle in CIELAB color wheel.

The amount of reward *R* (in number of liquid drops) that a target was worth depended on the distance between its color *θ* (in radian) and the template in the color wheel. The value function was a von-mises (normalized circular gaussian) of fixed concentration κ that was set to 2.5:

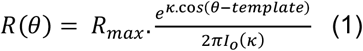

Where *I*_*o*_(κ) is the modified Bessel function of the first kind of order 0, such that the distribution sums to unity:

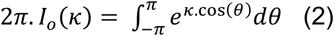

*R* was a number of drops and was rounded to the nearest unit. The maximum of reward *R*_*max*_ was fixed for monkey S (6 drops) and increased on each block for monkey B (12 drops + 1 drop for each completed block). The template color was fixed during a block such that subjects could learn it by trial and error, by choosing a color, receiving a feedback (number of drops) and updating the expected value for each color. The template color was randomly selected at the start of each block. To increase the likelihood of sampling from all colors during a session, we temporarily penalized the colors near already selected colors during the selection process such that they were less likely to be selected again.

At the start of each trial, the monkeys fixated a central cross on the screen. After a delay of 350±50ms, three colored disks appeared in three out of four possible locations (45, 135, 225, and 315 degrees from vertical). The colors were separated by at least π/6 rad on the color wheel. Monkeys had 2000ms to select a target by making a saccade to it and maintaining fixation for at least 50ms. After 100ms, monkeys received the number of drops associated with the chosen target. Only the chosen target stayed on screen while the reward was delivered. After 350/500ms for monkey B/S, a new trial started. If monkeys broke fixation before the onset of the targets or if they did not make a direct saccade to a stimulus, the screen changed color and a time-out of 200/500ms for monkey B/S was added to the inter-trial interval. In 20% of trials, the size of one of the targets was increased or decreased. This was random and did not predict reward or target location.

The template color was constant for a block of trials, allowing the monkey to learn and use the attentional template. The block (and the template color) would change when the monkey chose the target with the highest reward (i.e., the ‘best’ target) on 80/85% of 30 consecutive trials for monkey B/S. These template switches were uncued; the monkey could only detect the change through reward feedback. Blocks lasted between 80 and 575 trials (monkey B: 69 blocks, 138.2 ± 56.8 trials per block, monkey S: 102 blocks, 199.4 ± 114.1 trials per block). This was longer than the time it took monkeys to learn the template because it includes trials when the monkey broke fixation or did not make a direct saccade. To ensure a certain number of trials for data analysis, all blocks had at least 35 attempted trials before switching.

### Behavioral model

We modelled the animal’s choice using a Q-learning model with function approximation^37^. As in standard reinforcement learning models, the subject’s goal is to estimate the expected value *v* of each color *θ* in order to choose the best available color on each trial. Here, the state space is very large (100 colors) but continuous and structured according to the color wheel, so subjects can generalize what they learn about a color to the nearby colors (those that are closest in the color wheel). Hence, the expected value function can be represented not as a table (a value for each color) where each color is independent from the others, but as a parametrized function of N weight vectors with N being much less than the number of colors. In our case, the state space is represented by N equally distant radial basis functions *x*_*i*_ that are von-mises functions of concentration κ and centered on *μ*_*i*_ (with *μ*_1_ = 0 rad):

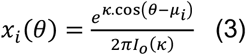

Where *I*_*o*_ is described in equation 2. The parameter κ controls the degree of generalization during learning, the smaller κ, the more generalization. The expected value function *v* at trial *n* is a linear function of the weights *w*_*i*_ and the radial basis functions *x*_*i*_:

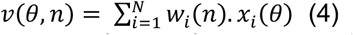

The weights were initialized at 0 at the start of the session. After receiving an amount of reward *R* for choosing the color *θ*_*chosen*_, all the weights *w*_*i*_ are updated according the learning rate *α*, the reward prediction error (RPE) and the distance between their centers *μ*_*i*_ and *θ*_*chosen*_:

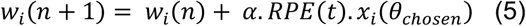

Where:

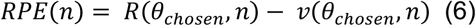

In this frame, the expected value function can be estimated for all colors at each trial.

We tested two variants of this model. In the ‘no reset’ model, the update follows equation 5, and the uncued template switches are not detected. In the ‘reset’ model, the weights *w*_*i*_ can be reset to 0 before doing the update. This occurred after a surprising event, when the absolute value of the RPE is above a threshold. The reset fraction *k*_*reset*_ was:

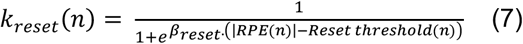

Where *β*_*reset*_ was set to 10^4^ such that *k*_*reset*_ was 0 or 1. Consecutive resets were penalized by increasing the *Reset threshold* based on the number of trials since the last reset (*# trials since last reset*). The smaller the *volatility* parameter, the greater the penalization:

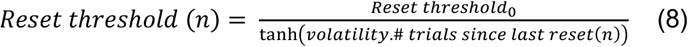

For this second model the full update rule was:

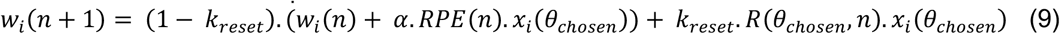

At each trial *n*, we computed the expected value *EV* of each option *θ* by adding potential biases for the location of the option (*Location*(*l*) = 1 at the location of the option and 0 otherwise), whether the size of the option was bigger or smaller than the standard size (*Size*(*s*) = 1 if the option was smaller (s=1) or bigger (s=2) and 0 otherwise), the proximity to a preferred colored *θ*_*pref*_ and the proximity to the previously chosen color *θ*_*prev chosen*_:

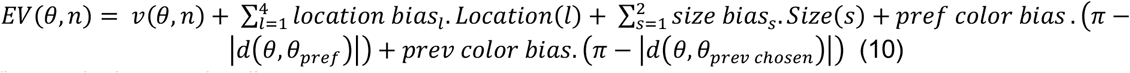

Where *d* is the angular distance:

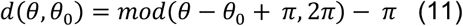

The choice probability *P* for each target *θ*_*j*_ was computed using the softmax rule:

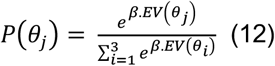

*β* was set to 0.3 for both monkeys.

### Behavioral model fit and selection

The ‘no reset’ model had 10 parameters and the ‘reset’ model had 12 parameters. Parameters were first fit using the Bayesian Adaptive Search (BADS) toolbox to get a first estimate of the parameters (https://github.com/lacerbi/bads). These parameters were used for the initialization of the fit with the Variational Bayes Monte-Carlo (VBMC) toolbox (https://github.com/lacerbi/vbmc)^81^. VBMC is an approximate inference method to fit and evaluate computational models. Specifically, it computes the approximate lower bound of the log model evidence (or log Bayes factor) for model comparison. We fit the models by first initializing the parameters using the BADS estimates (see Table 1). If the solution did not converge then we performed additional runs that were initialized using the posterior estimated on the previous run. All solutions converged.

**Table 1.**
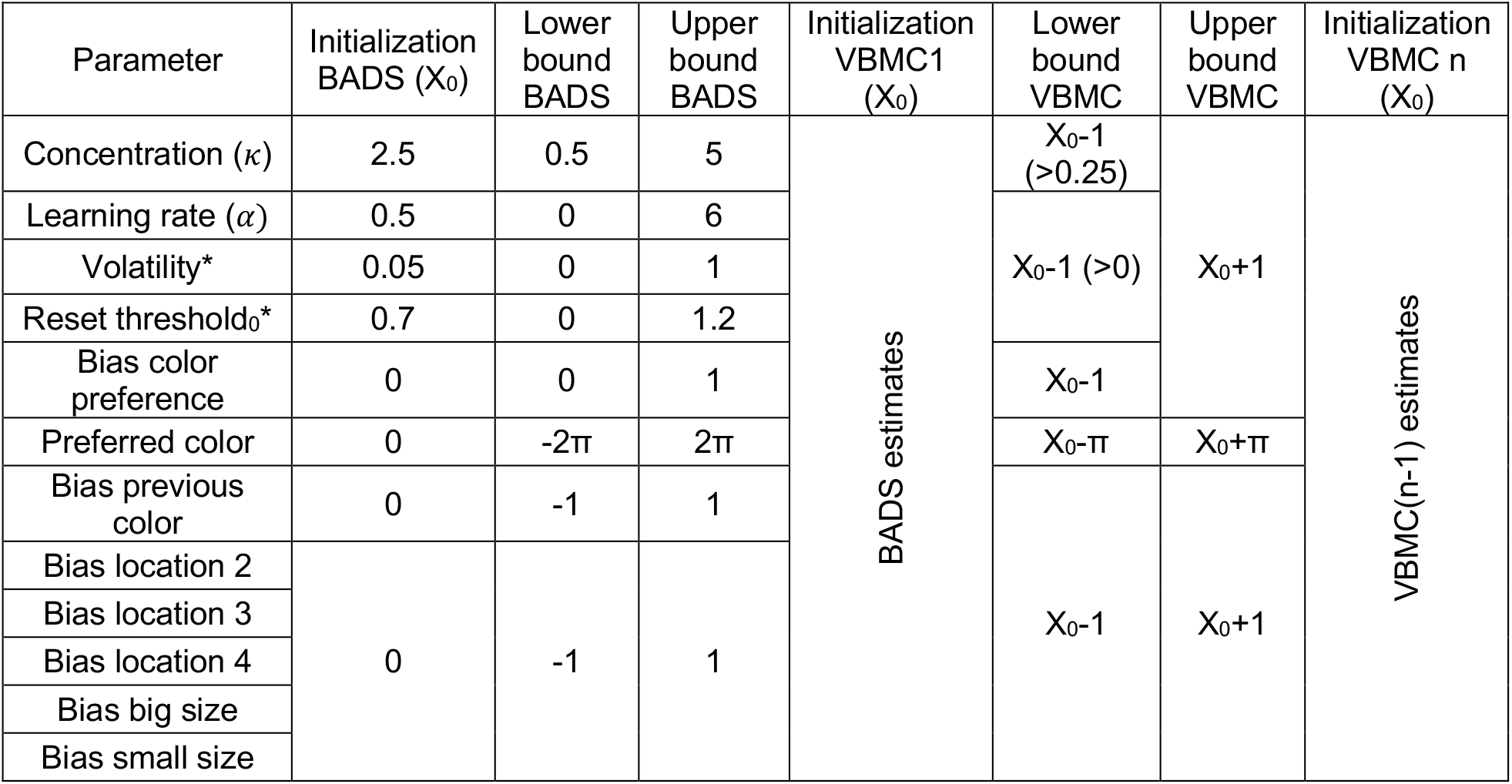
Priors for the difference steps of the behavioral model fit. *Reset model only.

**Table 2.** Mean and standard deviation of the parameters estimates for 6 channels. For the preferred color, we computed the parameter estimate as the mean angular distance and the standard deviation using the angular distance to the mean (equation 11).

| Parameter | Monkey B | Monkey S |
| --- | --- | --- |
| Concentration ($\kappa$) | 1.6411 ± 0.0216 | 1.8154 ± 0.0331 |
| Learning rate ($\alpha$) | 0.4858 ± 0.0103 | 0.2832 ± 0.0077 |
| Volatility | 0.5121 ± 0.2322 | 0.406 ± 0.0020 |
| Reset threshold<sub>0</sub> | 0.8751 ± 0.0013 | 0.6979 ± 0.0081 |
| Bias color preference | 0.0554 ± 0.0157 | 0.1956 ± 0.0082 |
| Preferred color (rad) | -1.6218 ± 0.2767 | -2.7138 ± 0.0899 |
| Bias previous color | 0.4183 ± 0.0170 | 0.4619 ± 0.0094 |
| Bias location 1 (fixed) | 0 | 0 |
| Bias location 2 | 0.1236 ± 0.0095 | 0.0574 ± 0.0069 |
| Bias location 3 | 0.0557 ± 0.0126 | 0.0207 ± 0.0069 |
| Bias location 4 | -0.0216 ± 0.0103 | 0.0811 ± 0.0070 |
| Bias big size | 0.1342 ± 0.0241 | -0.0487 ± 0.0456 |
| Bias small size | -0.8133 ± 0.0430 | 0.1 ± 0.0402 |

Parameters were estimated by sampling the posterior distribution 3.10^5^ times.

We fit a series of models with N varying from 3 to 8 radial basis functions and compared them using the Bayesian Information Criterion (BIC) and the lower bound of the log model evidence. Both statistics reached an asymptote for N=6 for both monkeys (see Fig. S1F) so this model was selected. Allowing for resets in the template improved the model’s accuracy (monkey B: ΔBIC =467, N=9,536 trials, monkey S: ΔBIC =406, N=20,338 trials; with 6 radial basis functions; Fig. S1F) and so resets were included in the model. Unfortunately, the number of resets was too few to reliably identify reset related neural responses.

On each trial, the ‘estimated template’ was computed as the color with highest value:

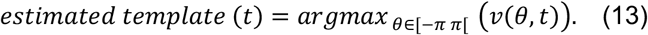

The entropy of the template was computed by transforming the value function in a strictly positive probability distribution:

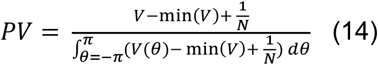

Where *V* is the value function and *N* is the number of possible colors (values of *θ*, 100).

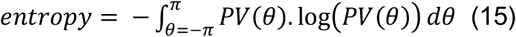

The model with two learning rates, one for positive and one for negative reward prediction errors, was an extension of model with a reset and so it had 13 parameters. It was fitted as described above with 6 radial basis functions. Model with one and two learning rates were compared using BIC.

### Electrophysiological recordings and signal processing

Animals were implanted with a titanium headpost to immobilize the head and with two titanium chambers to provide access to the brain. The chambers were positioned using 3D models of the brain and skull obtained from structural MRI scans. Chambers were placed to allow for electrophysiological recording from LPFC, FEF (frontal chamber) and LIP (parietal chamber).

Epoxy coded tungsten electrodes (FHC Inc, Bowdoin, ME) were used for both recording and micro-stimulation. Electrodes were lowered using a custom-built micro-drive assembly that lowered electrodes in pairs from a single screw. Recordings were acute; up to 60 electrodes were lowered through intact dura at the beginning of each recording session and allowed to settle for 2-3 hours before recording. This enabled stable isolation of single units over the session. Broadband activity (sampling frequency = 30 kHz) was recorded from each electrode (Blackrock Microsystems, Salt Lake City, UT). We performed 8 recording sessions in Monkey B and 9 sessions in Monkey S.

Based on MRI scans, we identified electrodes that were likely to be located in the FEF for each monkey and confirmed our identification using electrical micro-stimulation. Based on previous work, we defined FEF sites as those for which electrical stimulation elicited a saccadic eye movement^82^. Electrical stimulation was delivered in 200ms trains of anodal-leading bi-phasic pulses with a width of 400μs and an inter-pulse frequency of 330Hz. For monkey B, we found 4 sites that responded to electrical stimulation by evoking a saccade with a stereotyped eye movement vector at ≤100μA (reliability of the stimulation was: 83% at 50 μA, 67% at 50μA, 50% at 100μA, and 25% at 100μA). For monkey S, we found 5 sites that responded to electrical stimulation (reliability of the stimulation was 67% at 50μA, 57% at 50μA 28% at 50μA, 43% at 100μA, 67% at 100μA). Sites anterior and lateral to the FEF were labelled as LPFC.

Before the start of each session, monkeys performed a delayed match to sample task during which a target was presented for 125±25ms in 8 possible locations around a central fixation cross. After a delay, monkeys were instructed to saccade to the location of the sample. At the start of the session, this delay was short but rapidly increased to reach up to 650ms for monkey B and 600ms for monkey S. We identified pairs of electrodes located in LIP using the following criterion. We filtered the signal using a 4-pole 300-3000Hz band-pass Butterworth filter. We set the threshold for the detection of a spike by multiplying an estimate of the standard deviation of the background noise σ_*n*_ by a factor *β* that we varied between 1.8 and 5^83^:

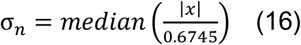

Time points at which the signal *x* crossed this threshold with a negative slope were identified as putative spiking events. Repeated threshold crossings within 32 samples (1.0667ms) were excluded. A channel was considered to be in LIP if there was a significant modulation of the spike rate during the delay (calculated by counting the total number of spikes during the delay) by the location of the remembered location of the sample in correct trials. We selected the value of *β* (= 2.2) that maximized the number of LIP channels detected using this method. Both electrodes in a pair were considered in LIP if at least one electrode reached this criterion.

Electrophysiological signals for the main task were filtered offline using a 4-pole 300Hz high-pass Butterworth filter. The spike detection threshold for all recordings was set equal to −4σ_n_ (equation 16). Repeated threshold crossings within 32 samples (1.0667ms) were excluded. Waveforms around each putative spike time were extracted and were manually sorted into single units, multi-unit activity, or noise using Plexon Offline Sorter (Plexon). The experimenter was blind to trials during all steps of preprocessing and spike sorting.

### Single neuron sensitivity to the factors of the task and model selection

To estimate the sensitivity of single neurons to the factors of interest, we used a 10-fold cross validation procedure (Fig. S2B). We fit the z-scored firing rates with the model representing the factors of interest using the Matlab function ‘fmincon’ with the default algorithm ‘interior-point’ that enables to solve a constrained minimization problem. To understand how neurons encoded task-relevant information, we fit four competing models for each individual neuron:

*Estimated template model*:

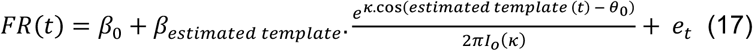

Where *I*_*o*_ is defined in equation 2, *estimated template* is defined in equation 13 and *e*_*t*_ is the residual. We used the same model for the true template model (Fig. S2F-G).

*Expected value model*:

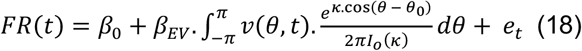

Where *v*(*θ, n*) is the value of the option color *θ* at trial *n* defined at equation 4.

*Mean value model* (equivalent to equation 18 with a κ close to 0):

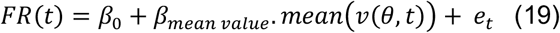

*Chosen color model*:

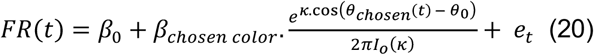

We used the following constraints: *β*_0_ ∈ [−10 10], *β*_*model*_ ∈ [−10 10], *θ*_0_ ∈ [−2*π* 2*π*] and κ = 10^*ρ*^ with *ρ* ∈ [−2.5 2.5] and checked that all solutions converged. We computed the explained variance (R^2^) of the model on the validation trials. We only included neurons with more than 500 trials in our analysis. A neuron was considered significantly sensitive to the factor of interest in the model if the mean R^2^ across validation sets was positive. Models were mutually exclusive such that if a neuron was significantly sensitive to several factors, the winning model was the one with the highest mean R^2^.

To estimate the significance of the proportion of neurons for each mutually exclusive model, we shuffled the firing rates of neurons (500 shuffles) and computed the proportion of significant neurons for each model. We then compared the proportion of neurons for each model to this null distribution.

### Pseudo-population decoding and cross-template generalization

To understand how the neural population represented the task, we combined neurons recorded during different sessions to create 100 independent pseudo-populations of N neurons for our regions of interest. N was the minimum number of neurons across regions that had at least 60 trials for each condition (some populations had more). For each pseudo-population, the activity matrix was a concatenation of the activity of N neurons (randomly selected with replacement) in m randomly selected trials in each condition. N and m were the same for all regions and across all time points and locations (when relevant) but specific to each analysis (estimated template, choice, chosen value, target color and chosen color).

For the estimated template (Fig. 2C-E), the true template (Fig. S2G) and stimulus color (Fig. S5B) analyses, we trained three classifiers: one for each color bin to discriminate the pseudo-population response at each time point, such that each classifier hyperplane separated the ‘in color bin’ vs. ‘out of color bin’ (with a randomly subsampled equal number of trials for each of the two other bins). For the estimated template and true template analyses, trials were balanced across progression in the block (1/3, 2/3, 3/3). For the stimulus color analysis, trials were balanced across chosen and unchosen stimuli (Fig. S5B). For the choice (Fig. 5E and S5D-E), chosen color (Figs. 5D and S4C) and chosen value (Figs. 5B and S5A), we trained one classifier. For the choice and chosen value analyses, trials were balanced across estimated template color with three bins. For the chosen color analysis, trials were balanced across estimated template color but with only two bins. In all cases, a subset (20%) of trials were withheld for validation, again balanced across conditions.

We used a standard linear support vector machine (SVM) classifier fit with the Matlab function ‘fitcsvm’ with the default solver ‘Sequential Minimal Optimization’. To minimize overfitting of the classifiers, fitcsvm uses the L1 norm for regularization (with box constraint as hyperparameter) and a kernel scaling parameter. The hyperparameters were set to minimize the 10-fold cross-validation loss. We used the default optimizer (‘bayesopt’) and the acquisition function ‘expected-improvement-plus’ that prevents the over-exploitation of one region of the parameter landscape. The projection of the trial input (the vector of the firing rates of the N neurons) onto the hyperplane defined the degree to which this trial is in or out of the class also called the score. We transformed this score in the posterior probability to belong to the class using the Matlab function ‘fitPosterior’.

The estimated template (Fig. 2C-E), the true template (Fig. S2G), and stimulus color (Fig. S5B) analyses used three classifiers and so we used two ways of calculating the overall classification accuracy. First, we took the class with the maximum posterior probability as the class for a particular trial. Second, we trained a feedforward, fully connected neural network for classification (function ‘fitcnet’ in Matlab with default settings) on the posterior probability across all three classes (withholding validation trials). The latter method tended to improve the classification accuracy and could account for distortions in the representation of the color wheel, so we report the results of this method. Classification accuracy was taken as the probability for a trial to be correctly classified on validation trials (chance level was 1/3). For all other analysis, that had only one classifier, we used the direct output of the SVM classification. Here as well, classification accuracy was taken as accuracy on validation trials (chance level was 1/2).

Statistical significance was estimated with one-sided z-tests against chance level (1/3 or 1/2 for one classifier) with a Bonferroni correction for multiple comparison for time series.

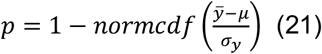

Where 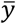 is the sample mean, σ_*y*_ is its standard deviation, *μ* is the hypothesized population mean and ‘normcdf’ is the function that returns the cumulative distribution of the standard normal distribution (computed with the Matlab function ‘normcdf’). Note that we used a z-test rather than a t-test because the sample size is defined arbitrarily by the number of bootstraps.

Cross-template generalization was assessed by training the SVM classifier using trials from one color bin of the estimated template and computing the classification accuracy either on validation trials from the same estimated template bin (‘within’) or from the other template bins (‘across’) (Fig. 5 and S5). Statistical significance was estimated with paired one-sided z-tests (‘within-across’) against 0 using the mean and standard deviation of the bootstrap distributions (equation 21) and with a Bonferroni correction for multiple comparison for time series.

### Principal Component Analysis

For the Principal Component Analysis (PCA; Fig. 2E and S2D), trials were sorted into 3 expected color bins and 20 time points (200ms windows centered on the onset of the targets), yielding 60 total conditions. The activity matrix X was the mean population activity (across trials) for each condition and each neuron. The principal components of this matrix were identified by decomposing the covariance matrix C of X using the Matlab function ‘pca’:

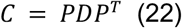

where each column of P is an eigenvector and D is a diagonal matrix of corresponding eigenvalues of C. The first 4 eigenvectors (in decreasing order of variance explained) respectively explained 95% / 89% / 90% of the variance in LIP/FEF/LPFC. We constructed a reduced 3-dimensional space using the first, second and fourth eigenvectors (Fig. 2E) or the first three eigenvectors (Fig. S2D) and projected the population activity vector for a given condition into this reduced dimensionality space. The percent explained variance and effective dimensionality were estimated with 100 bootstraps, by drawing with replacement from the 146 / 216 / 475 neurons in LIP/FEF/LPFC. The effective dimensionality was computed as:

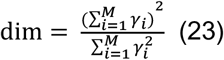

With M=59 as the degrees of freedom.

### Trial-by-trial estimated template decoding using population decoding

We used a decoding approach to estimate the trial-by-trial neural representation of the estimated template. These analyses were done independently for each session, on simultaneously recorded neurons. For each block, we isolated the first 35 trials of the block and labelled them as validation trials that we did not use to train the classifiers nor the neural network. We trained three classifiers, one for each color bin, on balanced trials across color bins as above. But here, we trained a feedforward, fully connected neural network (function ‘feedforwardnet’ in Matlab) with a hidden layer of size 10, the activation function *relu* (‘poslin’ in Matlab) which used as inputs the posterior probability to belong to each class for all trials except the validation trials and with an output layer of size 2 with a linear activation function that predicted the cosine and sine of the estimated template. Decoding accuracy was calculated on validation trials by calculating the circular distance (equation 11) between the estimated template (from the behavioral model) and neurally predicted estimated template.

The update was computed such that it should always be positive if it is i) toward the chosen color for a positive RPE and ii) away from the chosen color for a negative RPE:

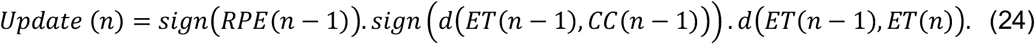

Where *d* is the angular distance (equation 11) and sign is the function that takes the value 1 if its argument is positive and -1 if it is negative. *ET* is the estimated template (either from the behavioral model or decoded from the neural population) and *CC* is the chosen color.

Circular correlations were computed using the MATLAB circular statistics (RRID:SCR_016651) toolbox.

### Generalized multiple linear regressions and representation alignments

For the generalized linear regressions used in Figs. 5, 6 and S6, we used the Matlab function ‘fitglm’. To remove any carry-over effect from previous trial, we removed the baseline firing rate (taken between - 600ms and -300ms from targets onset) from the firing rate in each 200ms window. The firing rates were z-scored for each window.

For the ‘global’ regression, we used all trials (Figs. 5, 6, and S6). The regressors were the reward magnitude, the value of the chosen stimulus, and the value of the unchosen stimulus with the highest value, both on the current trial and the previous trial. We also included the mean value of the current trial (taken as the mean value across all presented stimuli). For the ‘local’ value, the regressors were whether the stimulus was chosen, its value on chosen trials (0 otherwise), and its value on unchosen trials (0 otherwise). We only considered neurons where more than 500 trials were recorded, and with a firing rate that had enough variance to fit all the regressors.

At each time point, we defined the representation of a task factor (reward, chosen value and unchosen value) as the vector of the regression weights for this factor across neurons. We concatenated the distribution for the contralateral options (top right and bottom right on the screen) and the ipsilateral options (top left and bottom left on the screen) such that the distribution at each time point was 2 x N, with N as the number of neurons. The alignment *r* of these representations was taken as the Pearson correlation between the vectors (calculated using the Matlab function ‘corr’):

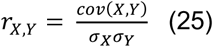

Where *cov* is the covariance and σ is the standard deviation. X and Y are the regressors estimated for each neuron on two different halves of the session. We performed 10 random splits of our sessions and estimated the mean correlation between the representations between the two sessions halves. To estimate the variability, we ran 5000 bootstraps, sampling N (global) or 2 x N neurons (local, the dimension of the vector, as above a concatenation of the two contralateral locations) with replacement.

We first estimated the split-half reliability of the representations by comparing the representation of the same factors computed in the two halves of the session (Figs. 5C, 6A-D and S6A-E). This gives us access to the maximal alignment that could be found between the representations. Statistical significance was estimated using a paired one-sided z-test against 0 with a Bonferroni correction for multiple comparisons across time. Comparison between strengths of reliability was estimated with a paired two-tailed z-test against 0.

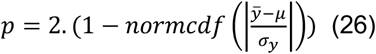

With the same variables as defined in equation 21.

When looking at the alignment between two different representations (either two different regressors or the same regressors at different locations), we only compared the representation in two different halves of the session such that the same trials were never used to compute the compared representations (Fig. 6CD and S6EG). To estimate the degree of alignment between representations, we computed the correlation disattenuation^54^:

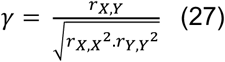

Where *r*_*X,Y*_ is the split-half correlation between the representation of the factors X and Y defined in equation 25, *r*_*X,X*_ and *r*_*Y,Y*_ are the split-half reliability of the representations X and Y respectively defined above. Statistical dissimilarly was estimated using a one-sided z-test against 0 with a Bonferroni correction for multiple comparisons across time points (22 time points).

To estimate the overall alignment across locations, we computed the area under the curve of the representation reliability (distance index = 0) or alignment across locations over time (22 time points, Fig. S6E) and over bootstraps (Figs. 6C and S6G). Statistical significance was estimated using a paired one-sided z-test (assuming that smaller distances would be more similar) against 0 with a Bonferroni correction for multiple comparisons (6 pairs).

To compute the z-scored explained variance of the chosen value representation across locations we used the split-half procedure described above. We estimated the chosen value regressor in one half of the session (either for the same of another location) and computed the R^2^ of the regression in the other half of the session. We performed 250 shuffles of the chosen value across trials and z-scored the R^2^ with the shuffled R^2^ distribution for each neuron. Statistical significance was estimated using a one-sided z-test against 0 across neurons. To estimate the overall explained variance across locations (Fig. S6F), we computed the area under the curve of the explained variance across time (22 time points, Fig. S6H) across neurons. Statistical significance was estimated using a paired one-sided z-test (assuming that smaller distances would be more similar) against 0 with a Bonferroni correction for multiple comparisons (6 pairs).

The peak of the unchosen value representation was computed by finding the time point at which the unchosen value reliability was maximal for each bootstrap. Statistical comparison between brain regions was estimated using a two-sample two-tailed z-test.

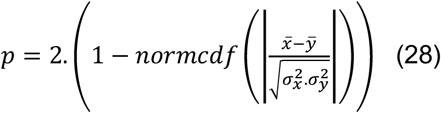

With the same variables as defined in equation 21.

#### Abbreviations

FEF: Frontal Eye Field
LIP: Lateral Intraparietal cortex
LPFC: lateral Prefrontal Cortex
RPE: Reward Prediction Error
+RPE: Positive Reward Prediction Error
-RPE: Negative Reward Prediction Error

## Supplemental information

**Fig. S1.**
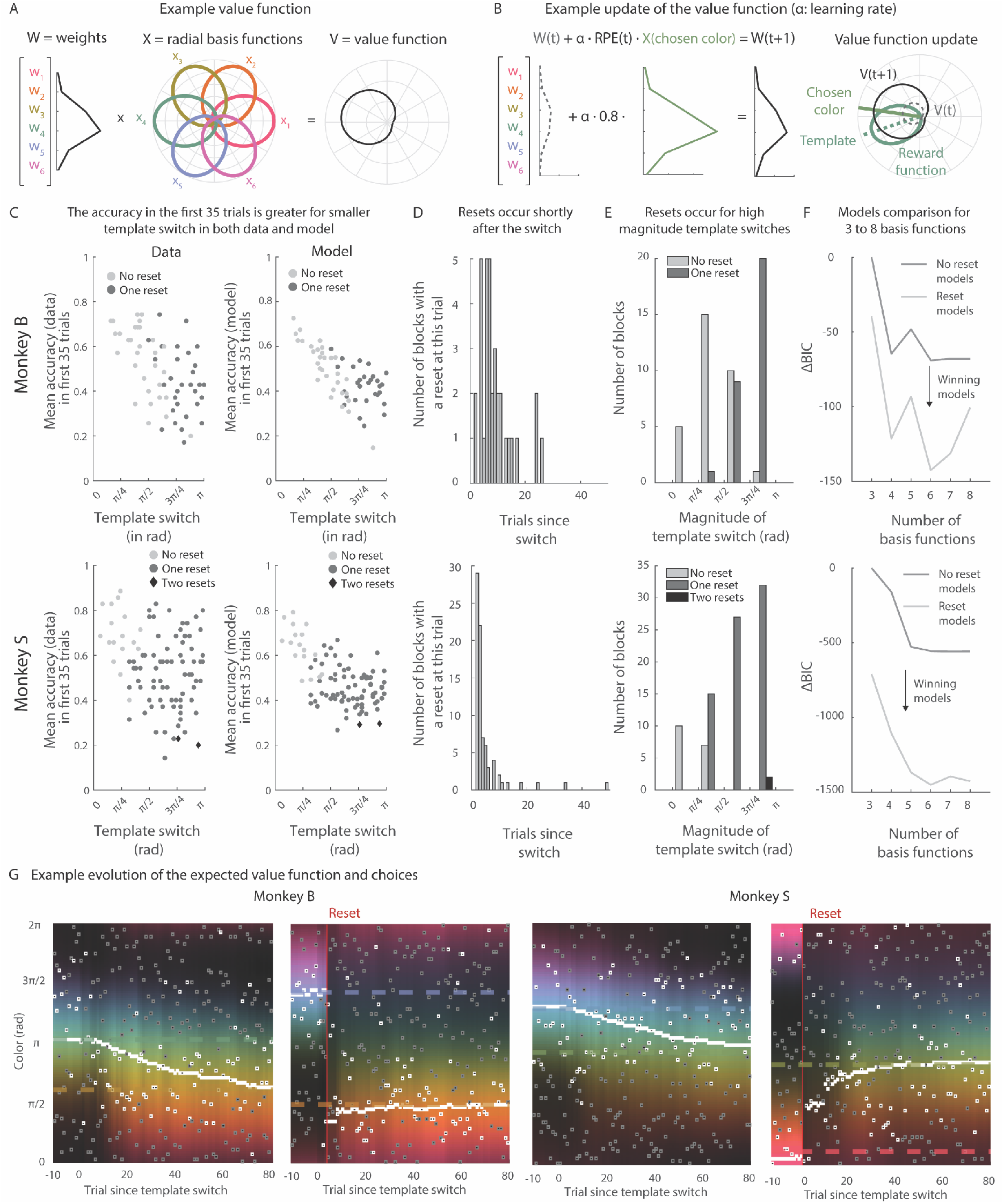
Extended figure1. (**A**) Example value function. On each trial, the value function is computed by combining the weights (left) and the radial basis functions (middle) centered on equally distanced colors to calculate the value function across colors (right). (**B**) Example update of the value function. After a positive reward prediction error (RPE), here 0.8 (a very large RPE), the model weights on the radial basis functions increase as a function of their distance to the chosen color (here green). This leads to an increased updated value function for colors closer to the chosen color. (**C**) Mean accuracy in the first 35 trials after a template switch for the subjects (left) and the model (right). The accuracy is greater when the previous and current templates are closer. Dot color and shape indicates whether a switch was detected in the model, this tends to occur for larger template switch. The model also captured the relationship between the magnitude of the change of template color and the monkey’s performance (r = -0.5455, p<0.001 for the data and r = -0.6943, p<0.001 for the model, 61 template switches / r = -0.2744, p=0.0078 for the data and r = -0.5740, p<0.001, 93 template switches, for monkey B/S) (**D**) Histogram of when resets are detected after the template switch for monkey B (top) and monkey S (bottom). (**E**) Histogram of how many resets were detected per bin of magnitude of template switch (absolute value of the angular distance between the previous and current templates). (**F**) Bayesian Information Criterion (BIC) of the ‘reset’ and ‘no reset’ models for 3 to 8 radial basis functions. (**G**) Examples of evolution of the expected value function (as in Fig 1G), chosen colors (white open squares) and unchosen colors (grey open squares) on each trial. The estimated template is indicated with a white full marker on each trial. The brighter the color, the higher the expected value according to the model. Previous and current templates are indicated with dashed lines. Chosen colors cluster around high values close to the estimated templates.

**Fig. S2.**
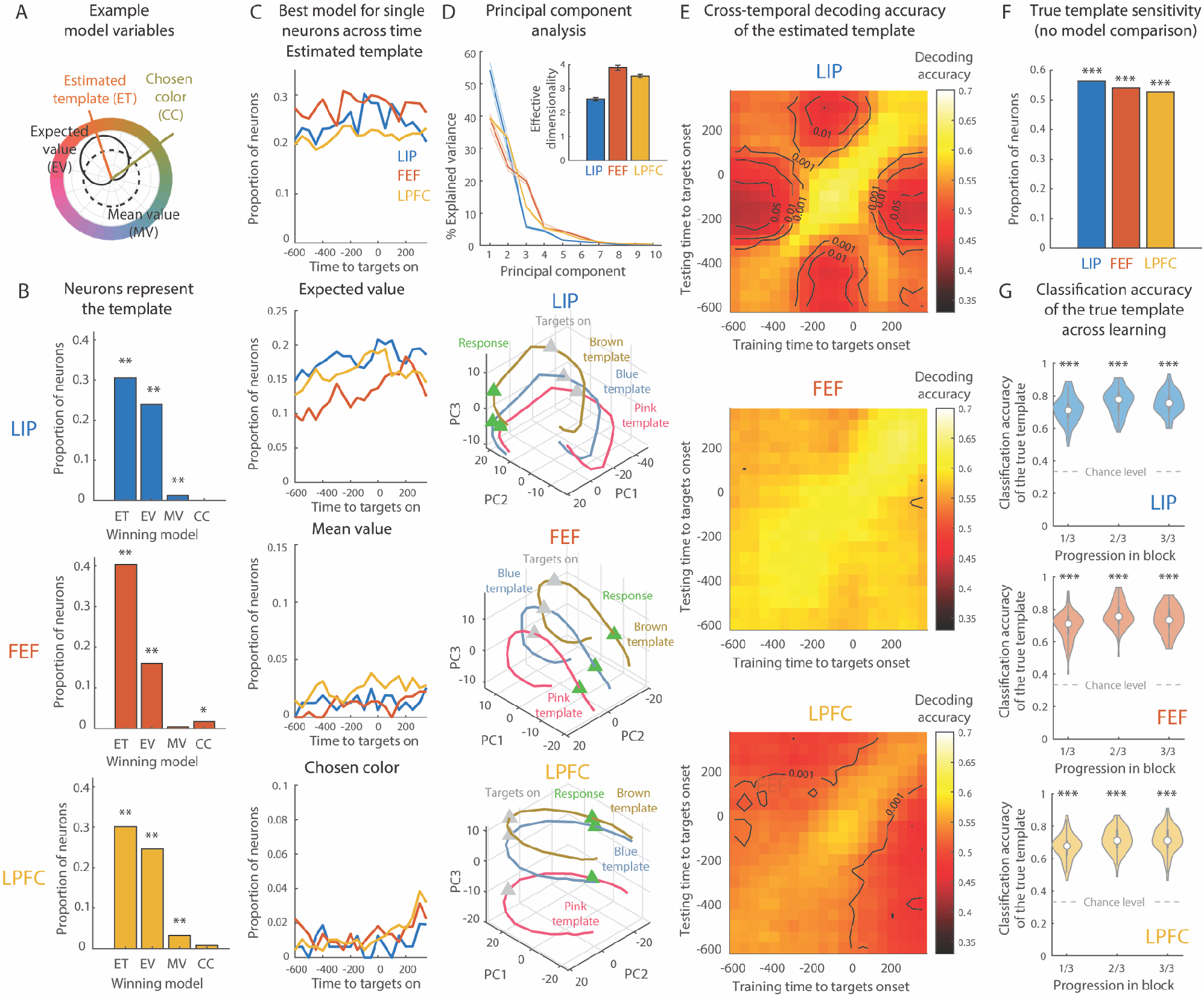
Extended figure 2. (**A**) Example trial showing the difference between the estimated template (orange), the chosen color (green), the expected value function (black) and the mean value (dashed black). (**B**) Proportion of significant neurons for the 4 mutually exclusive models: estimated template (ET), expected value (EV), mean value (MV) and chosen color (CC). Models were fit to 167/231/492 neurons in LIP/FEF/LPFC in a -600 to 300ms window around the onset of the targets. The expected template best explained neural activity in the largest group of neurons in all three regions (30.54/40.26/30.08% in LIP/FEF/LPFC, all p≤0.002, permutation test), although the expected value distribution was also well represented (29.95/16.02/24.59% of neurons in LIP/FEF/LPFC, all p≤0.002). Mean value and chosen color were comparatively less represented in all three regions (MV: 1.20/0.92/3.25% in LIP/FEF/PFC, p=0.008/0.0978/0.002 and CC: 0/1.73/0.23% in LIP/FEF/PFC, p=0.6447/0.01/0.1138). See methods for details on model comparison. (**C**) Proportion of neurons significantly encoding the four mutually exclusive models in each region from the onset of the targets (300ms windows, as in Fig. 2D). (**D**) Top panel shows percent explained variance by the first 10 principal components (100 bootstraps, mean and 95% confidence interval). Inset shows the effective dimensionality (100 bootstraps, mean and standard error to the mean). Bottom panels show neural activity in all three regions projected into a reduced dimensionality space consisting of the first three eigenvectors (in decreasing order of explained variance). LIP: 146 neurons, FEF: 216 neurons, LPFC: 475 neurons. The grey triangle represents the onset of the targets and the green triangle the approximate time of response (200ms after the onset of the targets). (**E**) Cross-temporal decoding of the attentional template (as in Fig. 2F, trained across all progression levels). X-axis corresponds to time window used to train classifier and y-axis to the time window used for testing the classifier (on withheld trials). Color code indicates classification accuracy of template and lines indicate significance level (z-test on 100 bootstraps). (**F**) Proportion of neurons significantly sensitive to the true attentional template (i.e., the color used by the behavioral task). Thus, the representation of the template did not depend on the behavioral model. (**G**) True template classification accuracy for each third of the block, estimated on withheld trials (as in Fig. 2E left half, 113 neurons per region, trained on 76 trials per progression level (three levels) and estimated template color bin (three bins)). Violin plot: central white dot is the median, thick vertical grey bar represents the 25^th^ to 75^th^ quartile and area represents the kernel density estimate of the data. For all panels, * p≤0.05, ** p≤0.01 *** p≤0.001.

**Fig. S3.**
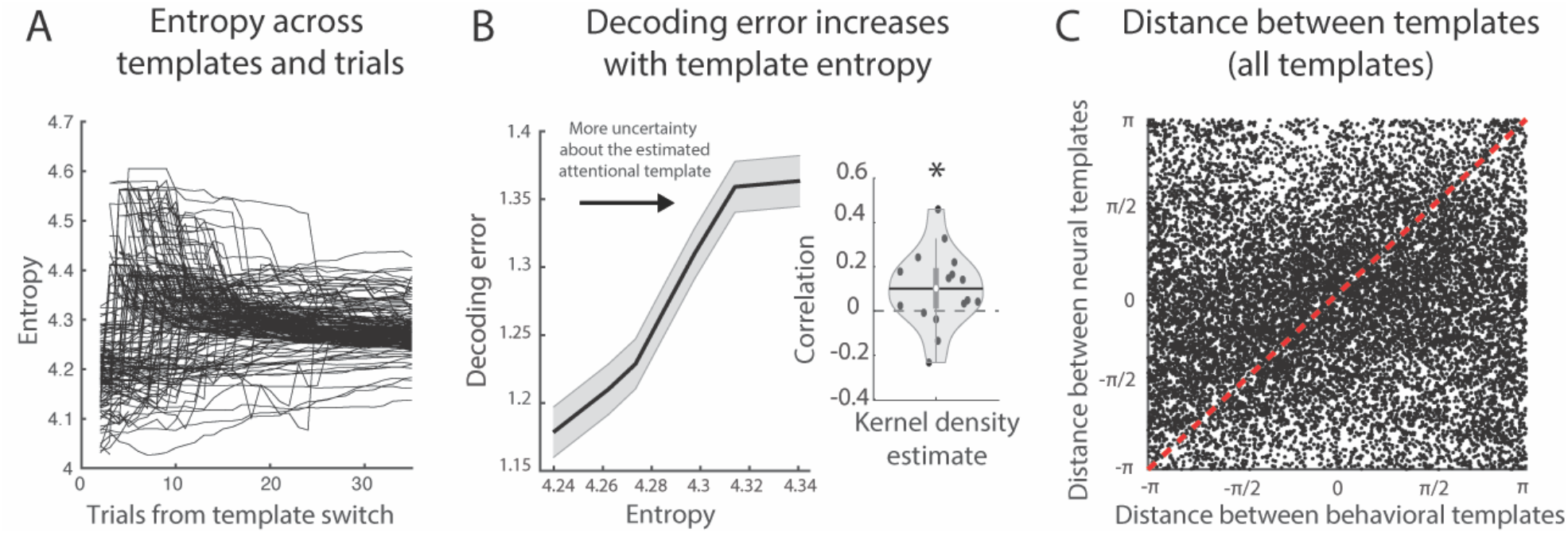
Extended figure. (**A**) Entropy of attentional template over trials, relative to template switch. Each line is a series of trials after the template switch (171 blocks). Entropy is often low before monkeys realize the template has changed. (**B**) Decoding error measured as absolute circular distance between neural and behavioral estimated templates (± SEM) for each bin of entropy (see methods, 10 bins, smoothing of 4 bins, all withheld trials). Violin plot shows mean Pearson correlation between the decoding error and the template entropy across validation trials for each session. The greater the attentional entropy, and therefore the uncertainty about the estimated template, the greater the decoding error (r(5,838)=0.0954, p<0.001, one-sided Pearson correlation). This effect was consistent across sessions (t(16)=2.4996, p=0.0118, one-sided t-test). (**C**) Scatter plot of circular distance between mean neural and behavioral estimated templates (r = 0.1092, p<0.001, 14,535 pairs, permutation test on the circular distance).

**Fig. S4.**
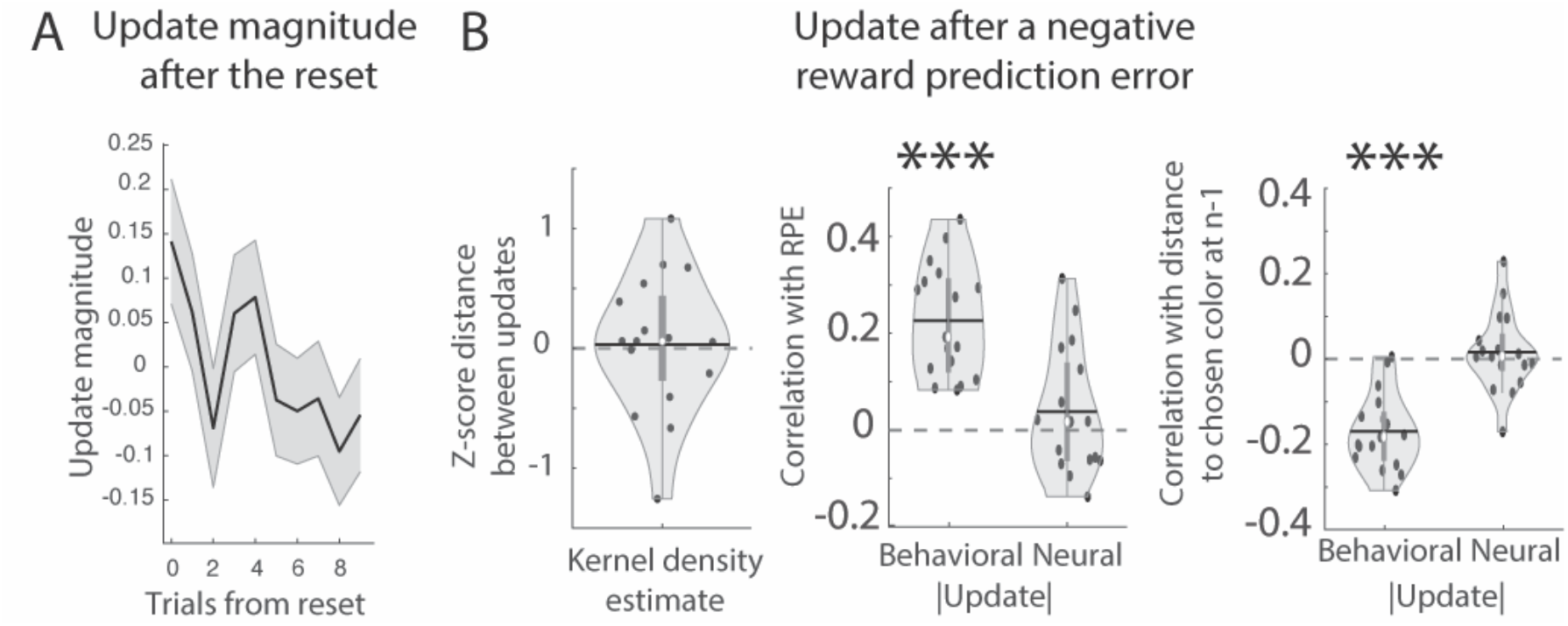
Extended figure 4. (**A**) Mean absolute angular distance between neural estimated template at trial n and n+1 over trials, relative to the reset (mean across blocks, ±SEM, N=111 blocks). The update magnitude decreased after the reset (GLM with factor trial after the reset, β=-0.019±0.007, t(1,108)=-2.6863, p=0.007). (**B**) *Left*. Mean z-scored distance between the neural and the behavioral update away from the chosen color on withheld trials, following a negative RPE. Black dots are individual sessions, central white dot is the median, horizontal black bar is the mean, thick vertical grey bar represents the 25^th^ to 75^th^ quartile and area represents the kernel density estimate of the data. *Middle*. Mean Pearson correlation between the update magnitude and the RPE magnitude on withheld trials. *Right*. Same as middle but for absolute distance between the estimated template before the update (at n-1) and the chosen color. For all panels, *** p≤0.001.

**Fig. S5.**
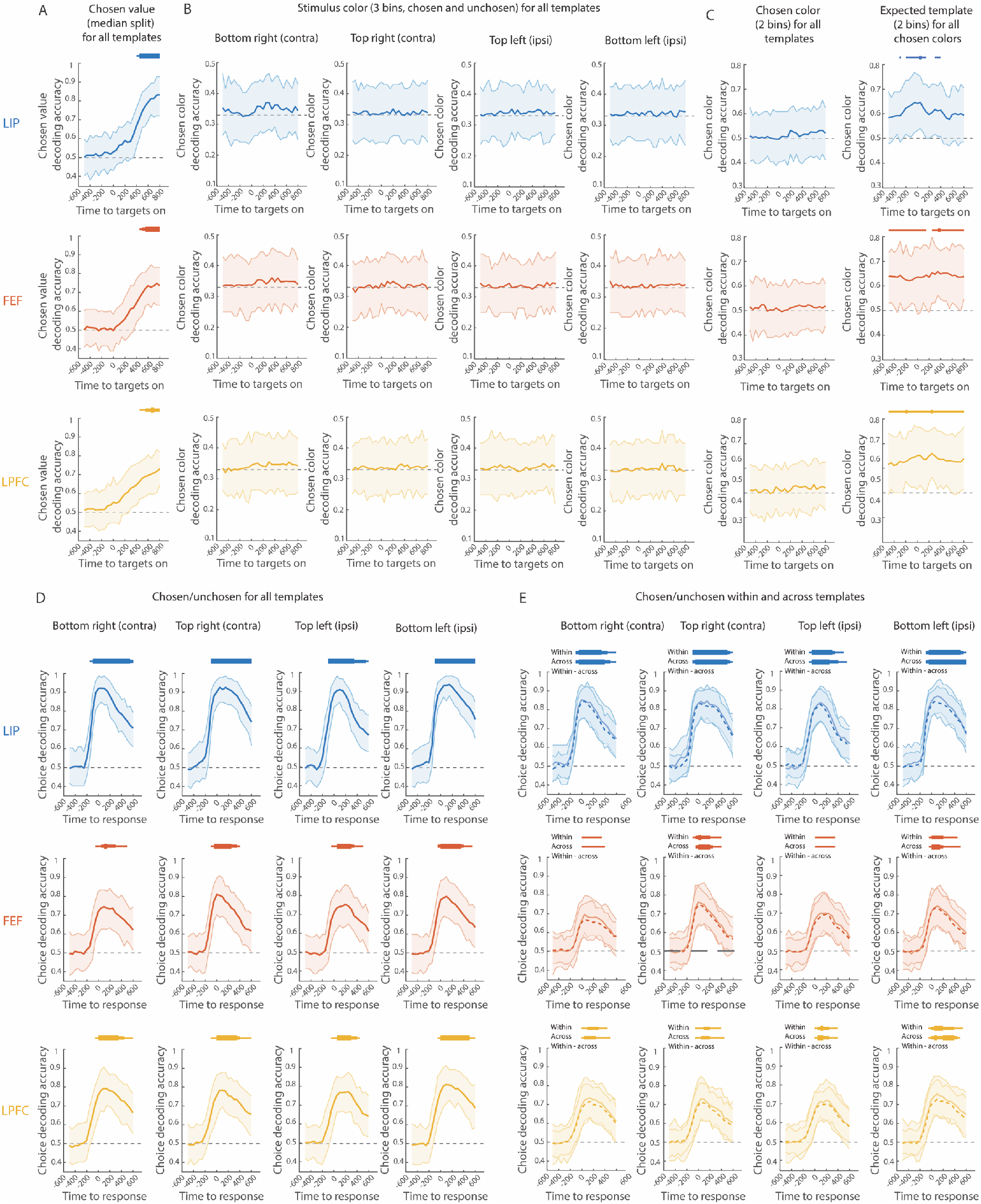
Extended figure 5. (**A**) Time course of classification accuracy of chosen value (with 95% confidence interval; 110 neurons, trained on 128 trials per estimated template bin, 3bins). Chance level was 1/2. For all panels, bar thickness indicates significance level: p≤0.01, p≤0.05 Bonferroni-corrected (27 time points) and p≤0.01 Bonferroni-corrected. (**B**) Time course of stimulus color classification accuracy of stimulus color (with 95% confidence interval). Computed on withheld trials for each time point (115 neurons, trained on 128 trials per stimulus color bin, 3bins; split between chosen and unchosen stimuli). Chance level was 1/3. p≥0.23 for all three regions. (**C**) Time course of classification accuracy of chosen color (*left*) and estimated template color (*right*; both with 95% confidence interval; 133 neurons, 80 trials per chosen color and estimated template bin, 2 bins for each). The chosen color could not be decoded when we balanced the estimated template color (all p≥0.28), while the estimated template could be decoded when we balanced the chosen color (p≤0.009 in all three regions, although this did not survive Bonferroni correction. Significance level threshold was lowered: bar thickness indicates significance level: p≤0.05, p≤0.01 and p≤0.05 Bonferroni corrected across time (27 time points) (**D)** Time course of classification accuracy of choice (with 95% confidence interval; on withheld trials; 90 neurons, trained on 120 trials per estimated template bin, 3bins). Chance level was1/2. Bar thickness indicates significance level: p≤0.01, p≤0.05 Bonferroni-corrected (23 time points and 4 locations) and p≤0.01 Bonferroni-corrected. (**E**) Same as D but computed on withheld trials with the same estimated template color as the training trials (solid line) or with a different estimated template color bin (dashed line).

**Fig. S6.**
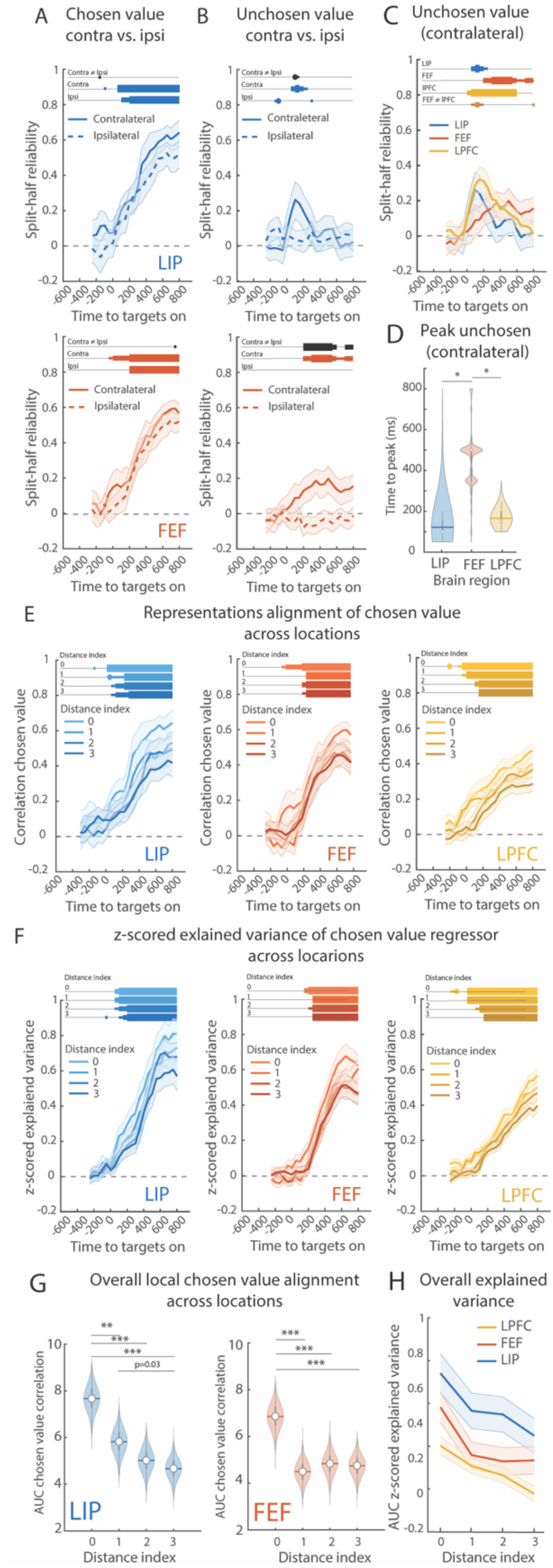
Extended figure 6. (**A**) Time course of the mean split-half reliability (with 95% confidence interval) of the chosen value regressors for the two contralateral locations (contra) and the two ipsilateral locations (ipsi, dashed line) for LIP (top) and FEF (bottom) (see Fig. 6A for LPFC). Bars indicate the significance of the split-half reliability (blue/red) and the difference in reliability strengths between the two regressors (black). (**B**) Same as A for the unchosen value for LIP and FEF (see Fig. 6B for LPFC). (**C**) Same as A for the unchosen value regressor in the contralateral locations for LIP, FEF and LPFC. Bars indicate the significance of the split-half reliability and the difference in reliability strengths between FEF and LPFC (orange). There was no difference between FEF and LIP (p≥0.0324) and LIP and LPFC (p≥0.0102). (**D**) Violin plot: time point at which the unchosen value reliability was maximal across bootstraps. Central white dot is the median, horizontal bar is the mean, thick vertical grey bar represents the 25^th^ to 75^th^ quartile and area represents the kernel density estimate of the data. Statistical significance was estimated using a two-sample two-tail z-test. * p≤0.05. (**E**) Time course of the mean split-half reliability (distance index = 0) and the correlation between chosen value regressors across locations (with 95% confidence interval) for LIP, FEF and LPFC (see methods for details). Bars indicate the significance of the split-half reliability (distance index = 0) and the correlation disattenuation between chosen value vectors. (**F**) Same as E but for the z-scored explained variance of the chosen value regressor (see methods for details). (**G**) Violin plot: mean area under the curve of the representation reliability (distance index = 0) or alignment across locations across time (22 time points, shown in E) across bootstrap for LIP an FEF. Central white dot is the median, horizontal bar is the mean, thick vertical grey bar represents the 25^th^ to 75^th^ quartile and area represents the kernel density estimate of the data. Statistical significance was estimated using a paired one-sided z-test (assuming that smaller distances would be more similar) using the reliability or the correlation disattenuation against 0. * p≤0.05, ** p≤0.01, *** p≤0.001 Bonferroni-corrected (6 pairs). (**H**) Mean area under the curve of the z-scored explained variance of the chosen value regressor (shown in F, see methods for details).

